# The role of the L421P mutation in Penicillin-Binding Protein 1 (PBP1) in the evolution of chromosomally mediated penicillin resistance in *Neisseria gonorrhoeae*

**DOI:** 10.1101/2025.06.27.662027

**Authors:** Gabriella Gentile, Bryan Guzman, Adriana Le Van, Ann E. Jerse, Yonatan H. Grad, Daniel Dominguez, Tatum D. Mortimer, Robert A. Nicholas

## Abstract

*ponA*^L421P^ encodes a mutated variant of penicillin-binding protein 1 (PBP1) and is a key resistance determinant that increases the penicillin MIC (MIC_PEN_) above the clinical breakpoint in *Neisseria gonorrhoeae*. Despite the removal of penicillin from treatment guidelines for gonococcal infections in the 1980s, *ponA*^L421P^ is present in nearly 50% of current *N. gonorrhoeae* isolates in the PubMLST database. Bioinformatic analysis indicates that *ponA*^L421P^ is exclusive to *N. gonorrhoeae* isolates, whereas Leu-421 is 100% conserved in other *Neisseria* species. To understand the involvement of *ponA*^L421P^ in antibiotic resistance, we introduced *ponA* variants encoding 16 different amino acids at position-421 into FA6140, a penicillin-resistant gonococcal isolate that naturally harbors *ponA*^L421P^. Proline-421 was the only mutation that increased the MIC_PEN_ to the same level as FA6140. We also assessed the fitness of strains with the 16 mutant *ponA* alleles over multiple serial passages, both with and without sub-MIC levels of penicillin. There was no fitness defect attributed to *ponA*^L421P^ under these experimental conditions; instead, our analyses suggest that the widespread occurrence of *ponA*^L421P^ is driven by its capacity to increase the MIC_pen_ above the clinical breakpoint. In FA6140 transformed with the mosaic *penA* allele from strain H041, a ceftriaxone-resistant isolate, *ponA*^L421P^ increased the MIC of ceftriaxone, suggesting that ceftriaxone targets PBP1 in this strain. We conclude that the *ponA*^L421P^ allele emerged in gonococcal isolates, increasing the MIC_PEN_ above the clinical breakpoint, and has remained in the population even after the removal of penicillin from treatment guidelines.

**Importance:** The emergence of antibiotic-resistant *Neisseria gonorrhoeae* threatens effective treatment of gonorrhea, one of the most common sexually transmitted infections worldwide. Understanding the genetic changes that drive and maintain resistance is crucial for anticipating future resistance trends. Here, we investigated the impact of a key resistance mutation in PBP1 (encoded by *ponA*^L421P^). Although penicillin has not been used to treat gonorrhea for decades, this mutation remains widespread even in recent *N. gonorrhoeae* isolates. *ponA*^L421P^ confers clinically relevant penicillin resistance without imposing an *in vitro* fitness cost. *ponA*^L421P^ also increases resistance to ceftriaxone in strains with *penA* alleles that are associated with ceftriaxone resistance. This work highlights the role of the *ponA*^L421P^ allele in shaping the current antibiotic resistance landscape and supports the need for ongoing surveillance and evolutionary studies of such mutations in the gonococcal population.

## Introduction

*Neisseria gonorrhoeae* is the etiologic agent of the sexually transmitted infection gonorrhea, which causes ∼80 million infections annually (1, 2). The β-lactam antibiotic penicillin was first used to treat gonorrhea in the early 1940s. For the following 40 years, penicillin remained effective, but the minimum inhibitory concentrations of penicillin (MIC_PEN_) steadily increased, requiring higher doses to cure infections (3). By 1985, over 5% of circulating *N. gonorrhoeae* strains had MIC_PEN_ values above the breakpoint (≥2 μg/ml), prompting a switch in treatment guidelines to other antibiotics. The rise in penicillin resistance was driven by two mechanisms: plasmid-mediated production of penicillinases (TEM-1 and TEM-135 β-lactamases) and expression of resistance determinants that are variants of chromosomal genes (4–11). In the US, the percentage of penicillinase-producing *N. gonorrhoeae* isolates (PPNG) decreased by the 1990s, while the percentage of chromosomally mediated penicillin-resistant *N. gonorrhoeae* isolates (CMRNG) increased (2, 8).

β-lactam antibiotics target penicillin-binding proteins (PBPs), which are involved in the cross-linking of the peptide chains of peptidoglycan. *N. gonorrhoeae* has four PBPs, two of which (PBP1 and PBP2) are essential. PBP1 is a bifunctional PBP that catalyzes both glycan polymerization (transglycosylation) and peptide cross-linking (transpeptidation), and PBP2 is a monofunctional PBP, catalyzing transpeptidation reactions. Because PBP2 has a much higher rate of acylation than PBP1 for penicillin and other β-lactam antibiotics used to treat *N. gonorrhoeae* in susceptible strains, PBP2 is considered the lethal target for these antibiotics. CMRNG is facilitated by the step-wise transfer (12–14) of at least four resistance determinants from a penicillin-resistant strain to an antibiotic-susceptible strain: 1) mutant *penA* alleles, which encode variants of PBP2 that reduce the acylation rate of penicillin by 16-fold (15); 2) *mtr* mutations, which increase *mtrCDE* operon expression and result in increased penicillin efflux through the MtrCDE efflux pump (9, 16); 3) mutations in *porB*, which encodes the major outer membrane porin, that reduce the influx of penicillin into the periplasm (7, 17); and 4) *ponA*^L421P^, encoding a PBP1 variant with an L421P missense mutation. Mutations in *penC* were thought to contribute to high-level penicillin resistance (8), but *penC* was later shown to encode the pore-forming secretion protein, PilQ (18). These *pilQ* mutations compromise its secretin function and increase the MIC of multiple antibiotics (18, 19), but to date, *pilQ* mutations have not been observed in clinical isolates, as they disrupt the type IV pili that play a key role in gonococcal pathogenesis (20, 21). While the first three resistance determinants are well characterized in the literature, the contribution of *ponA*^L421P^ to penicillin resistance remains less well understood.

Our laboratory has shown that the L421P mutation in PBP1 lowers the acylation rate by 2– to 3-fold for penicillin G and other β-lactam antibiotics (8), but it is not clear whether this lower acylation rate explains the role of *ponA*^L421P^ in facilitating penicillin resistance. Transformation of the *penA, mtr,* and *porB* determinants from strain FA6140, a high-level-penicillin-resistant clinical isolate (MIC_PEN_ = 4 µg/ml), into the antibiotic-susceptible strain FA19 (MIC_PEN_ = 0.01 μg/ml) results in MIC_PEN_ between 0.5 to 1.0 µg/ml (12, 13), but further transformation with FA6140 DNA or *ponA*^L421P^ does not increase the MIC_PEN_ (22). All attempts to reach donor levels of resistance in gonococcal isolates by natural transformation have been unsuccessful (8, 10, 12). By contrast, we showed that reversion of *ponA*^L421P^ in two penicillin-resistant clinical isolates, including FA6140, back to the wild-type *ponA* allele decreases the MIC_PEN_ 2-fold (22). These data imply that *ponA*^L421P^ requires a yet to be determined genetic background that cannot be transferred between strains by transformation to exert its phenotypic effect on resistance.

We hypothesized that *ponA*^L421P^ arose during the years of penicillin use and has continued to remain in the population because *ponA*^L421P^ continues to confer a fitness benefit, even with the current β-lactam antibiotics recommended for use. In the present study, we sought to test this prediction by investigating the origins of *ponA*^L421P^ to elucidate the relationship between *ponA*^L421P^ and the evolution of β-lactam resistance in *N. gonorrhoeae*.

## Results

### *ponA*^L421P^ is exclusive to *N. gonorrhoeae* among *Neisseria* species

*N. gonorrhoeae* isolates can acquire resistance-conferring mutations from commensal *Neisseria* species and from *N. meningitidis* (23–25). We examined whether *ponA*^L421P^ arose via transfer of DNA from other *Neisseria* species, or if this mutation is specific to gonococcal isolates. We accessed the sequences of all unique *ponA* alleles across *Neisseria* species from the PubMLST Database, and divided them into three groups: (1) *N. gonorrhoeae* (n = 20,513 isolates, 172 *ponA* alleles), (2) *N. meningitidis* (n = 75,612 isolates, 897 *ponA* alleles), and (3) all other commensal *Neisseria* species (n = 1,604 isolates, 369 *ponA* alleles).

The PBP1 protein sequences across all three groups of *Neisseria* isolates are highly conserved (Fig. S1), particularly the region surrounding residue 421 (Fig. 1). Leu-421 is 100% conserved across all *N. meningitidis* and commensal *Neisseria ponA* alleles (Fig. 1B-C), but for *N. gonorrhoeae* isolates, residue 421 is either a leucine or a proline (Fig. 1A). Moreover, *ponA*^L421P^ was observed in 49.5% of all gonococcal isolates in the PubMLST database and was not found in other *Neisseria* species (Table S5). These data strongly suggest that *ponA*^L421P^ arose in *N. gonorrhoeae* and not through horizontal gene transfer from other *Neisseria* species.

**FIG 1.**
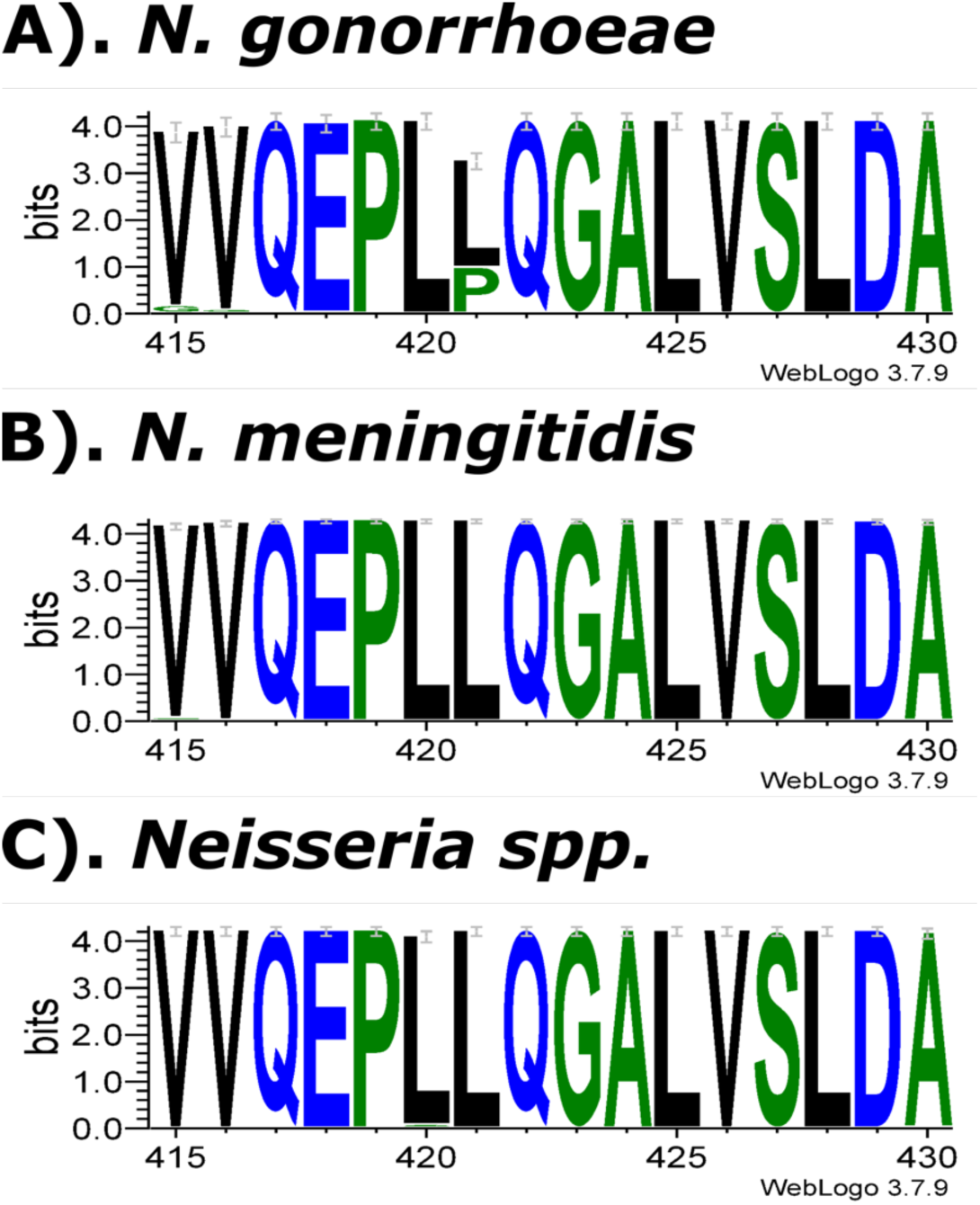
Sequence logo diagrams comparing the conservation of positions 415-430 in PBP1 from various *Neisseria* isolates. Diagram generated on WebLogo with MUSCLE-aligned *ponA* sequences. Sequences were downloaded from PubMLST and are a representative sub-set of all of the *ponA* alleles documented for each *Neisseria* species group: *N. gonorrhoeae* (n=172), *N. meningitidis* (n=897), and *Neisseria spp.* (n=369). The X-axis indicates the position of the amino acid. The height of the logo element (letter) represents its log-transformed frequency displayed in bits of information, and the overall height of the stack indicates the sequence conservation at that position.

### *ponA*^L421P^ is prevalent amongst gonococcal isolates with chromosomally-mediated penicillin resistance

To investigate the emergence and distribution of *ponA*^L421P^ in the gonococcal population, we assembled a dataset of sequenced *N. gonorrhoeae* isolates and the associated MIC_PEN_ values (n=16,891). The first appearance of *ponA*^L421P^ in a sequenced isolate was in 1961, and it rapidly expanded in the population in the 1980s (Fig. 2). Interestingly, we first observed the insertion of aspartate after position 345 in *penA* (Asp345a), which is a hallmark of early penicillin-resistant isolates (26, 27), in our dataset of sequenced isolates in 1956, only a few years prior to the L421P mutation in *ponA* (Fig. 2). We examined the phylogenetic distribution of *ponA* alleles in a representative sample of *N. gonorrhoeae* isolates without plasmid-mediated resistance (n=5,192) and found *ponA*^L421P^ emerged across multiple lineages in the gonococcal population (Fig. 3).

**FIG 2.**
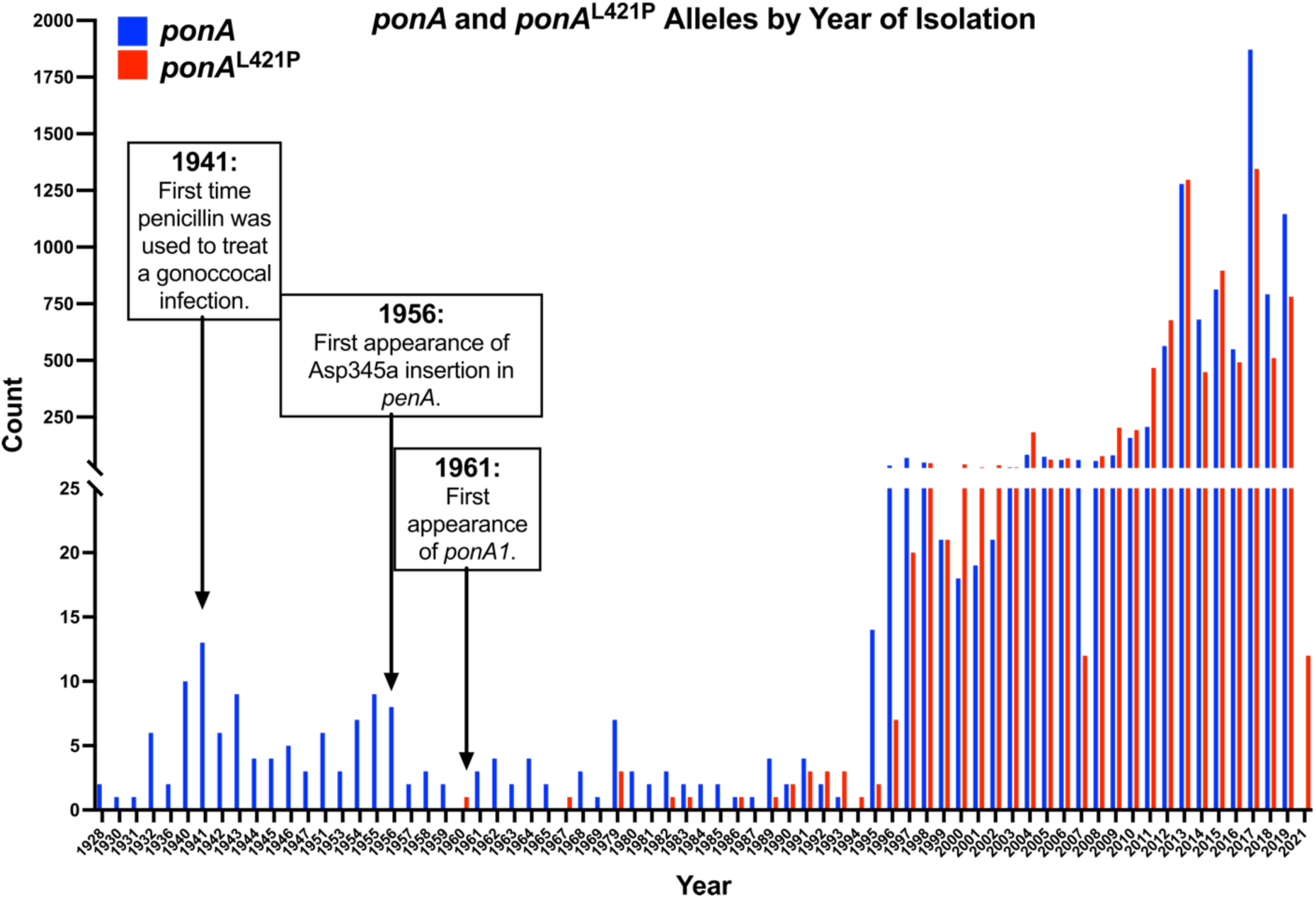
Historical analysis of *N. gonorrhoeae* isolates (n=16,891). Isolates are represented by the year the isolate was collected. Asp345a refers to the insertion of asparagine at position 345 in *penA* (encodes PBP2), a known penicillin-resistance-conferring mutation. Count represents the number of isolates that harbor *ponA* (blue) or *ponA*^L421P^ (red).

**FIG 3.**
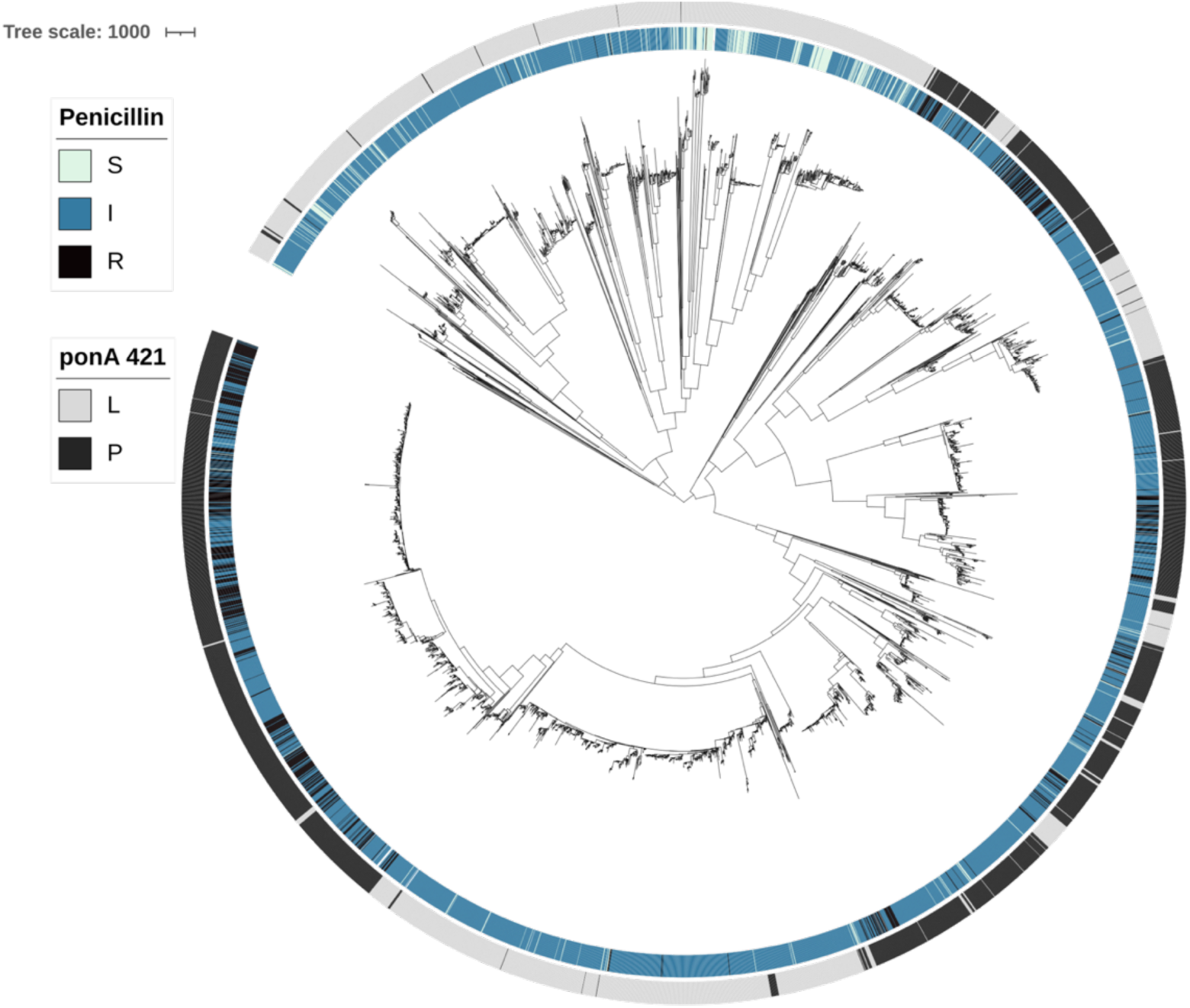
Phylogenetic distribution of *ponA* alleles in a representative sample of *N. gonorrhoeae* isolates without plasmid-mediated resistance (n=5,192). The outer annotation ring indicates the presence of *ponA* (L421, grey) and *ponA*^L421P^ (L421P, black). The inner ring indicates the level of penicillin resistance. S is susceptible (≤0.06 μg/mL), I is intermediate (0.25 – 1 μg/mL), and R is resistant (≥2 μg/mL) breakpoints of PenG as defined by the Clinical and Laboratory Standards Institute (CLSI) (83).

We next examined the relationship between *ponA*^L421P^ and penicillin-resistant CMRNG strains (Fig. 4, n=8,999). Less than 1% of isolates harboring the wild-type *ponA* allele included in Fig. 4 have resistant MIC_PEN_ values (≥2 μg/mL), while the majority (90%) fall within the intermediate MIC_PEN_ range (0.12–1 μg/mL). In contrast, 25% of isolates harboring *ponA*^L421P^ have resistant MIC_PEN_ (≥2 μg/mL), and 75% of isolates have intermediate MIC_PEN_. Few isolates, regardless of the *ponA* or *ponA*^L421P^ allele, had susceptible MIC_PEN._ Additionally, 91.61% of isolates that harbor mosaic *penA* alleles, which encode highly remodeled PBP2 variants that confer increased resistance to cephalosporins, also harbor *ponA*^L421P^ (Fig. 4B).

**FIG 4.**
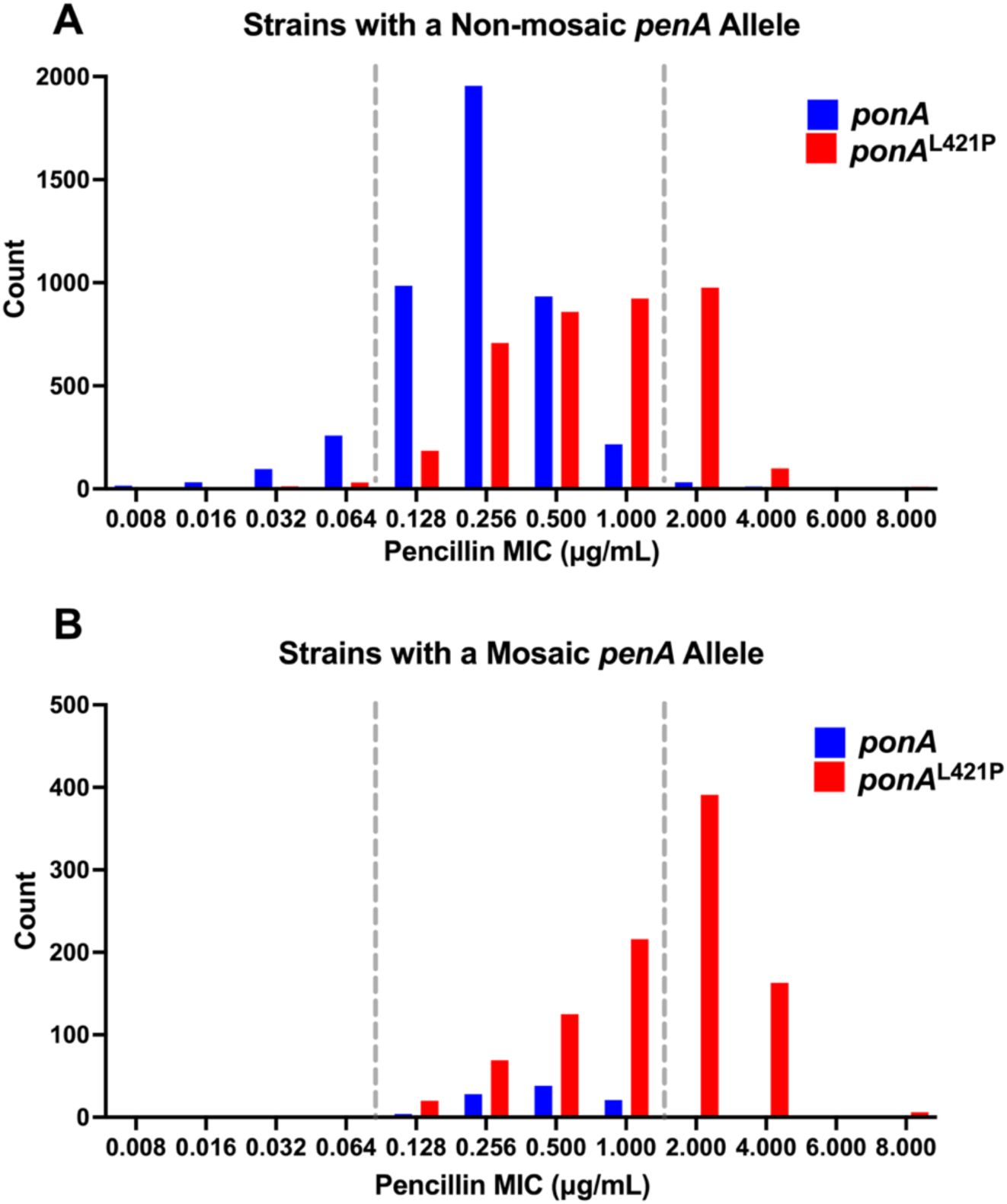
Distribution of MIC values for penicillin G (PenG) among *N. gonorrhoeae* isolates (n = 8,999). Strains are separated by harboring either (A) a non-mosaic *penA* allele, or (B) a mosaic *penA* allele. The X-axis shows the PenG MIC (MIC_PEN_), and dotted lines indicate the susceptible (≤0.06 μg/mL), intermediate (0.25 – 1 μg/mL), and resistant (≥2 μg/mL) breakpoints of PenG as defined by CLSI (83). The Y-axis shows the number of isolated *N. gonorrhoeae* strains for each MIC_PEN_, value. Red bars show distribution of strains with the *ponA*^L421P^ allele and blue bars shows the number of strains with the wild-type *ponA* (L421) allele. The mosaic *penA* alleles in isolates were defined and typed according to the NG-STAR database (56).

### Identification of other polymorphisms in gonococcal *ponA*

Amongst *N. gonorrhoeae* isolates in PubMLST, there were only two other mutations observed in gonococcal PBP1 at frequencies above 1%: A375T (7.07%) and Y537C (2.35%) (Fig. 5). In keeping with these frequencies, a phylogeny of 6,082 representative gonococcal genomes reveals these mutations are present in fewer lineages, compared to the phylogenetic distribution of the L421P mutation (Fig. S2). We next examined the relationship between each mutation and penicillin-resistant CMRNG strains (n=8,999). The majority (95%) of isolates harboring the A375T mutation in *ponA* have intermediate (0.12–1 μg/mL) to resistant (≥ 2 μg/mL) MIC_PEN._ This proportion does not change in isolates with alanine at residue 375 of *ponA*, as 95% of these isolates also have intermediate to resistant MIC_PEN_. The same trend is observed with the Y537C mutation. We do not observe A375T and Y537C in sequenced isolates collected during the penicillin treatment era, as they first appear in 1998 and 2001, respectively, well after penicillin was removed as a treatment for gonorrhea infections in the US in 1985. It remains unclear whether A375T and Y537C are directly involved in facilitating resistance to penicillin, but, given the fact that isolates with those mutations are not enriched for penicillin resistance and they are present in fewer gonococcal lineages than L421P, we did not interrogate them further.

**FIG 5.**
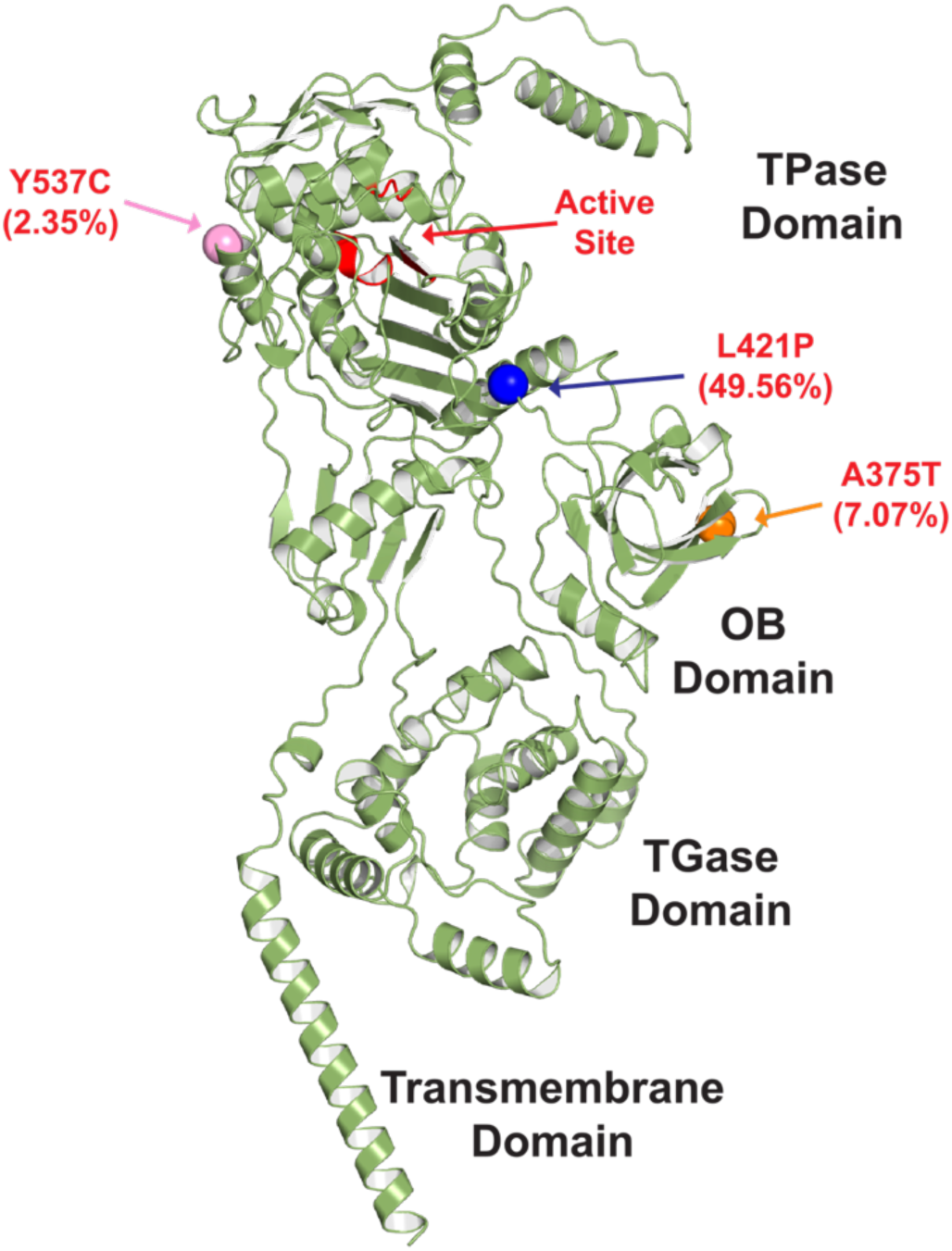
Polymorphisms of *N. gonorrhoeae* PBP1 mapped onto the predicted protein structure. Polymorphisms are colored according to their prevalence in the gonococcal population. Only polymorphisms detected at frequencies greater than 1% are shown: L421P (47.15%), A375T (7.07%), and Y537C (2.35%). The Transpeptidase (TPase), Transglycosylase (TGase), and the Oligosaccharide-Binding (OB) domains of PBP1 are labeled, with the TPase active site residues marked in red. MUSCLE-aligned sequences of gonococcal PBP1 were taken from PubMLST isolate database in order to identify polymorphisms and calculate population frequencies, see Materials and Methods. Predicted structure of gonococcal PBP1 is from the AlphaFold Protein Structure Database (AFDB).

### Only proline and serine at PBP1 position 421 increase the MIC_PEN_

To understand why only a proline mutation at residue 421 is observed in PBP1, we created an isogenic library of 16 mutant strains in FA6140, each with a different amino acid at position 421 (Table 1). FA6140 (MIC_PEN_= 4 μg/mL) naturally harbors *ponA*^L421P^, and reversion to *ponA*^421L^ reduces the MIC_PEN_ to 2 μg/mL (8, 22) (Table 1). In the strain library, FA6140 *ponA*^L421P^ was the only strain with an MIC_PEN_ of 4 μg/mL. FA6140 *ponA*^L421S^, which had a median MIC_PEN_ of 2.5 μg/mL, was the only other mutation that increased the MIC_PEN_ > 2 μg/mL, with all the other strains having the same MIC_PEN_ as FA6140 *ponA* (Table 1).

**Table 1.** FA6140 *ponA*^L421X^ Isogenic Mutant Library and penicillin MICs.

| Strains Name<sup>a</sup> | Amino Acid at Position 421 in *ponA* | Codon | Penicillin MIC (μg/mL)<sup>b</sup> |
| --- | --- | --- | --- |
| **FA6140<sup>c</sup>** | Proline (P) | CCG | **4** |
| **FA6140 *ponA*<sup>d</sup>** | Leucine (L) | CTG | **2** |
| **FA6140 *ponA*<sup>L421P</sup>** | Proline (P) | CCG | **4** |
| **FA6140 *ponA*<sup>L421L</sup>** | Leucine (L) | CTG | **2** |
| **FA6140 *ponA*<sup>L421A</sup>** | Alanine (A) | GCC | **2** |
| **FA6140 *ponA*<sup>L421R</sup>** | Arginine (R) | AGA | **2** |
| **FA6140 *ponA*<sup>L421C</sup>** | Cysteine (C) | TGC | **2** |
| **FA6140 *ponA*<sup>L421N</sup>** | Asparagine (N) | AAC | **2** |
| **FA6140 *ponA*<sup>L421E</sup>** | Glutamic Acid (E) | GAA | **2** |
| **FA6140 *ponA*<sup>L421G</sup>** | Glycine (G) | GGT | **2** |
| **FA6140 *ponA*<sup>L421H</sup>** | Histidine (H) | CAT | **2** |
| **FA6140 *ponA*<sup>L421Q</sup>** | Glutamine (Q) | CAG | **2** |
| **FA6140 *ponA*<sup>L421W</sup>** | Tryptophan (W) | TGG | **2** |
| **FA6140 *ponA*<sup>L421Y</sup>** | Tyrosine (Y) | TAT | **2** |
| **FA6140 *ponA*<sup>L421S</sup>** | Serine (S) | AGT | **2.5** |
| **FA6140 *ponA*<sup>L421F</sup>** | Phenylalanine (F) | TTC | **2** |
| **FA6140 *ponA*<sup>L421I</sup>** | Isoleucine (I) | ATC | **2** |
| **FA6140 *ponA*<sup>L421T</sup>** | Threonine (T) | ACT | **2** |

<sup>a</sup>All strains are created in the FA6140 background, a penicillin-resistant gonococcal isolate, as described in Methods and Materials. Each strain possesses a different amino acid at position 421 in *ponA*.
<sup>b</sup>The MICs reported are the median values determined from a minimum of three independent experiments.
<sup>c</sup>Penicillin-resistant gonococcal isolate, FA6140, which harbors *ponA*<sup>L421P</sup>.
<sup>d</sup>FA6140 with *ponA*<sup>L421P</sup> replaced with WT *ponA*.

### *In vitro* serial passaging of the FA6140 *ponA*^L421X^ mutant library in the absence or presence of sub-MIC concentrations of penicillin

The 2-fold higher MIC_PEN_ conferred by *ponA*^L421P^ and the timing of its first appearance in sequenced isolates suggest that *ponA*^L421P^ emerged in response to selection by penicillin. To test this, we performed *in vitro* serial passaging on our set of FA6140 *ponA*^L421X^ mutants in increasing sub-MIC_PEN_ concentrations (0.25, 0.5, and 1.0 μg/ml). Illumina sequencing revealed that equal amounts of each FA6140 *ponA*^L421X^ mutant were used to inoculate the cultures at hour 0 across all 3 trials (Fig. 6A, Fig. S3). In the absence of penicillin, the amino acids proline, cystine, asparagine, and glycine at position 421 appear to be more fit in this growth environment, as they have the largest fold increase in read counts at hour 36 relative to the inoculum (Fig. 6C). The *ponA*^L421P^ reads consistently doubled by hour 36 in all three trials, while the *ponA*^421L^ reads remained similar to, or decreased from, the number of reads in the inoculum. The proline and serine mutants were the only strains with an MIC_PEN_ above 2 μg/mL (Table 1), but the read counts for the serine mutant at hour 36 did not surpass its read counts at hour 0 (Fig. 6C). None of the strains differed significantly in growth rate from the parental strain (Table S4).

**FIG 6.**
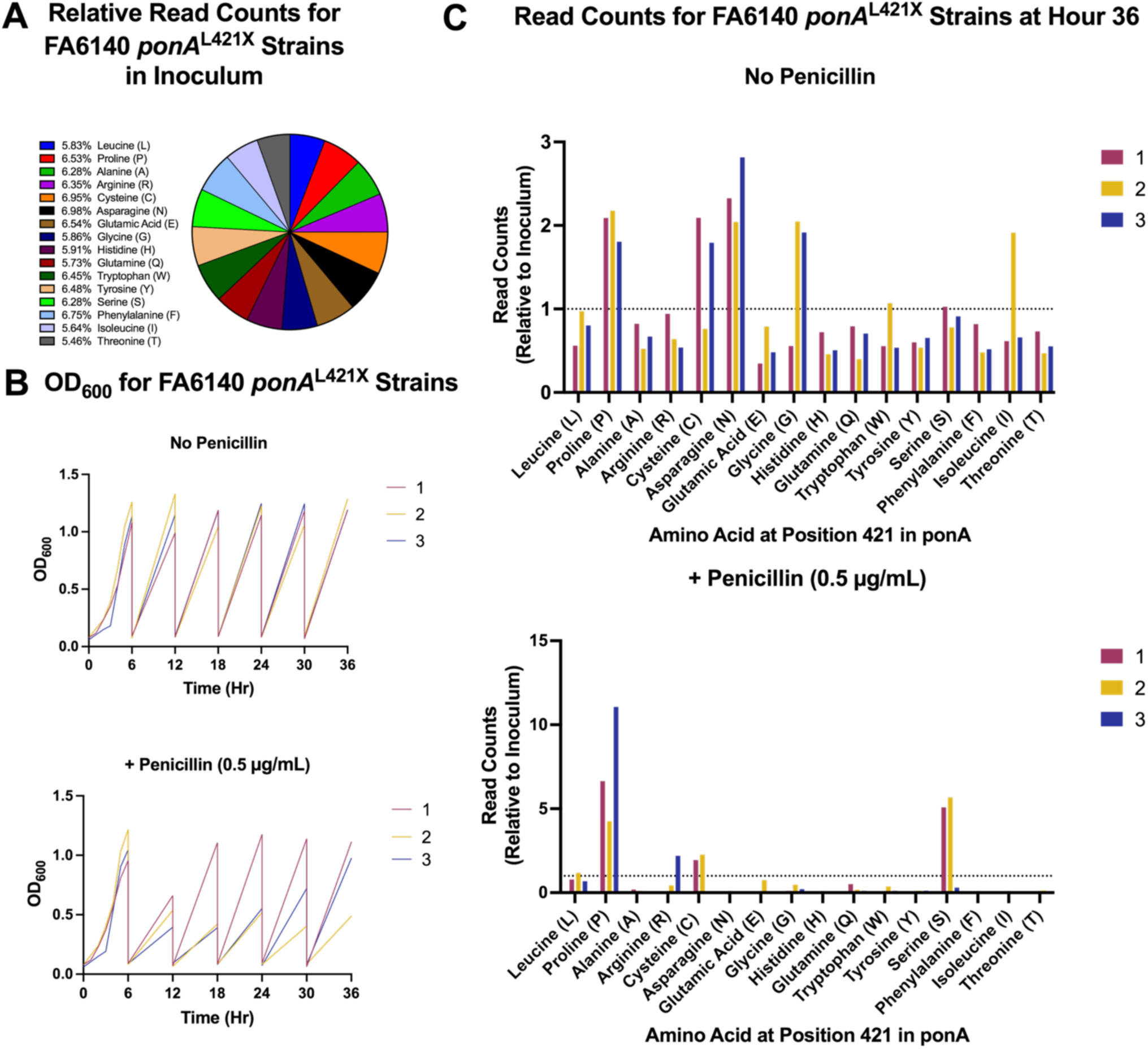
Optical densities (OD_600_) and Illumina read counts from *in vitro* serial passaging of the FA6140 *ponA*^L421X^ mutant library in the absence or presence of sub-MIC concentrations of penicillin. (A) Percentage of reads, relative to the total number of reads, for each mutant at hour 0 (inoculum). The inoculum was used to inoculate each culture at 0.08 OD_600_. (B) The serial passaging scheme included a total of 4 cultures, 3 of which were supplemented with different sub-MIC concentrations of penicillin (0.25, 0.5, and 1.0 µg/mL), and one with no penicillin added. Cultures were passaged every 6 hours, for a total of 36 hours, always with a starting OD_600_ of 0.08. (C) OD_600_ and (D) Illumina read counts of the cultures with no penicillin, compared to cultures supplemented with 0.5 µg/mL penicillin. (D) Reads for each mutant in the inoculum were normalized to 1, as indicated by the dotted line at y=1. Bars represent read counts for each mutant at hour 36, relative to their inoculum value. This experiment was performed 3 separate times (trials) and each trial represents a separate Illumina sequencing run; (C-D) Trial 1 is shown in red, trial 2 in yellow, and trial 3 in blue. The pie chart in (A) is a representative figure from trial 1, but all 3 trials had similar results. See supplemental for read counts from the cultures supplemented with 0.25 µg/mL of penicillin (Fig. S4) and for the inoculum used in trials 2 and 3 (Fig. S3).

We grew the collection of strains in growth medium with increasing concentrations of penicillin to examine which strains would grow in the presence of selective pressure. Optical Density at 600 nm (OD_600_) values during growth in medium containing 0.25 mg/ml penicillin were very similar to no antibiotic (Fig. S4), while there was more variability in the OD_600_ values across trials for the cultures supplemented with 0.5 μg/mL of penicillin (Fig. 6B). Of the 16 mutants, the two with the highest MIC_PEN_, FA6140 *ponA*^L421P^ and FA6140 *ponA*^L421S^, increased in read counts when grown in 0.5 μg/mL penicillin, while all the others decreased in read counts. FA6140 *ponA*^L421P^ consistently increased in read counts across all three trials, while read counts for FA6140 *ponA*^L421S^ increased in trials 1 and 2, but not in trial 3. In trial 1, FA6140 *ponA*^L421P^ had more read counts at 36 h than FA6140 *ponA*^L421S^, and both strains had similar reads counts at 36 h in trial 2 (Fig. 6C). FA6140 *ponA*^L421P^ is the only mutant that consistently increased in read counts over 36 h in cultures either with or without sub-MIC_PEN_ concentrations.

At the highest sub-MIC_PEN_ concentration tested, we observed more variable results. After 6 h of growth and passaging into fresh medium with 1 μg/ml penicillin, the OD_600_ increased very little over the next 12 h (Fig. 7A). However, in two of the three trials (1 & 3), there were still live cells present, as later passages showed measurable growth during the 6 h period. These data suggest that gonococcal cells remained viable but were not dividing during the prior 6 h growth periods and eventually acquired either phenotypic or genetic changes that allowed for measurable growth at the later growth periods. In trial 1, we observed increasing growth during consecutive passages, beginning at 18 h. Illumina sequencing of the *ponA* region revealed that serine was the predominant mutation at position 421 (Fig. 7B).

**FIG 7.**
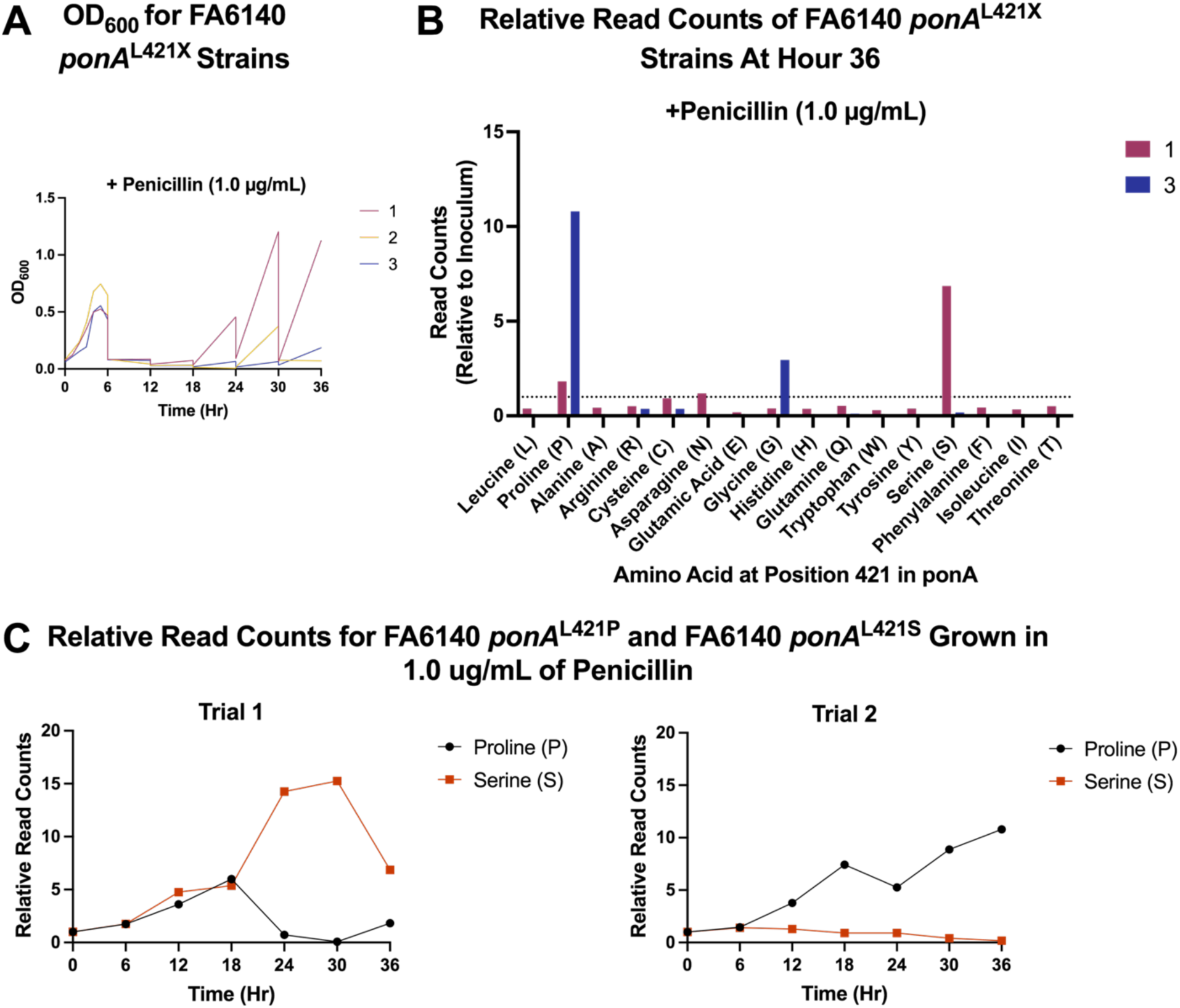
A *pilQ* mutation arose in the culture supplemented with 1.0 ug/ml of penicillin. Optical densities (A) and Illumina read counts (B) for the FA6140 *ponA*^L421X^ strains grown in cultures supplemented with 1.0 µg/mL of penicillin. (B) Read counts for each mutant at hour 36, relative to their inoculum (Hr 0) value. Inoculum reads for each mutant were normalized to 1, as indicated by the dotted line at y=1. (C) Read counts over time for FA6140 *ponA*^L421S^ (orange) FA6140 *ponA*^L421P^ (black), relative to their inoculum value (y=1), during trials 1 and 3. Each trial represents a separate Illumina sequencing run (A-B, trial 1 = red, trial 2 = yellow, trial 3 = blue). Cells growing in the culture from trial 2 appeared to die at hour 18 (A; yellow line), resulting in poor sequencing reads. See supplemental Fig. S5 for trial 2 data.

### Whole genome sequencing of cells from *in vitro* serial passaging FA6140 *ponA*^L421X^ mutant library reveals emergence of a mutation in *pilQ*

To determine if the FA6140 *ponA*^L421X^ mutants acquired additional mutations during the 36 hours of *in vitro* growth, we sequenced the genomes of cells collected at 0 h and 36 h. We looked for mutations that were distinct from the inoculum and present in greater than 20% of the reads, as anything detected below this would likely not be responsible for any changes in the overall codon counts from the Illumina reads (Figs. 6D, 7B). Across all 3 trials, most, but not all, mutations above the 20% threshold were either errors in the reference genome sequence used for alignment (not true genetic changes), also present in the inoculum, or in genes that would most likely be inconsequential to resistance.

In trial 1, we detected a mutation in *pilQ* at 36 h at 85% frequency in the culture supplemented with 1.0 μg/mL of penicillin. The mutation was an insertion of two bases (+CG) at position 459, which would result in a frameshift and early termination. Previous work from our lab showed that missense and frameshift mutations in *pilQ* increase MIC_PEN_ by inactivating PilQ and decreasing permeation of drug into the gonococcal cell (8, 18, 19, 28), which likely explains the marked increase in codon counts for the serine mutation at 24 h seen in trial 1 (Fig. 7C). Our data suggest that during trial 1, FA6140 *ponA*^L421S^ acquired a mutation in *pilQ* between hours 18 and 24, which allowed it to expand in the culture compared to FA6140 *ponA*^L421P^.

### *ponA*^L421P^ increases the ceftriaxone MICs of gonococcal isolates that also harbor a mosaic *penA* allele

Our data indicate that *ponA*^L421P^ is involved in penicillin resistance, but penicillin was removed as a recommended treatment in 1985, and ceftriaxone has been the only recommended β-lactam antibiotic since 2012 for treating gonococcal infections. Given that *ponA*^L421P^ appears to confer a modest fitness deficit *in vivo* (29), it might be expected that *ponA*^L421P^ would decrease in prevalence without selection by penicillin use. However, *ponA*^L421P^ is observed in a high percentage of isolates with a mosaic *penA* allele (91.61%, Fig. 5B), which is the predominant determinant for decreased susceptibility to ceftriaxone (30, 31). One possibility to explain the prevalence of *ponA*^L421P^ is that *ponA*^L421P^ influences the MIC of ceftriaxone in the presence of a mosaic *penA* allele.

To test this hypothesis, we replaced the *penA4* allele in FA6140 and FA6140 *ponA* strains with *penA41*, the mosaic allele from the multi-drug resistant gonococcal isolate, H041 (32). This allele encodes a PBP2 with a markedly reduced acylation rate for penicillin, cefixime, and ceftriaxone (31, 33, 34). Reversion of *ponA*^L421P^ to wild-type *ponA* reduced the MIC_PEN_ in both strain backgrounds, regardless of the *penA* allele (Fig. 8A), suggesting that PBP1 is the lethal target of penicillin in these strains. While reversion of *ponA*^L421P^ to *ponA* in strain FA6140 had no effect on the MIC of ceftriaxone (Fig. 8B), reversion of *ponA*^L421P^ to *ponA* in strain FA6140 *penA41* caused a 3-fold decrease in the MIC of ceftriaxone, suggesting again that PBP1 is the lethal target of ceftriaxone in this strain.

**FIG 8.**
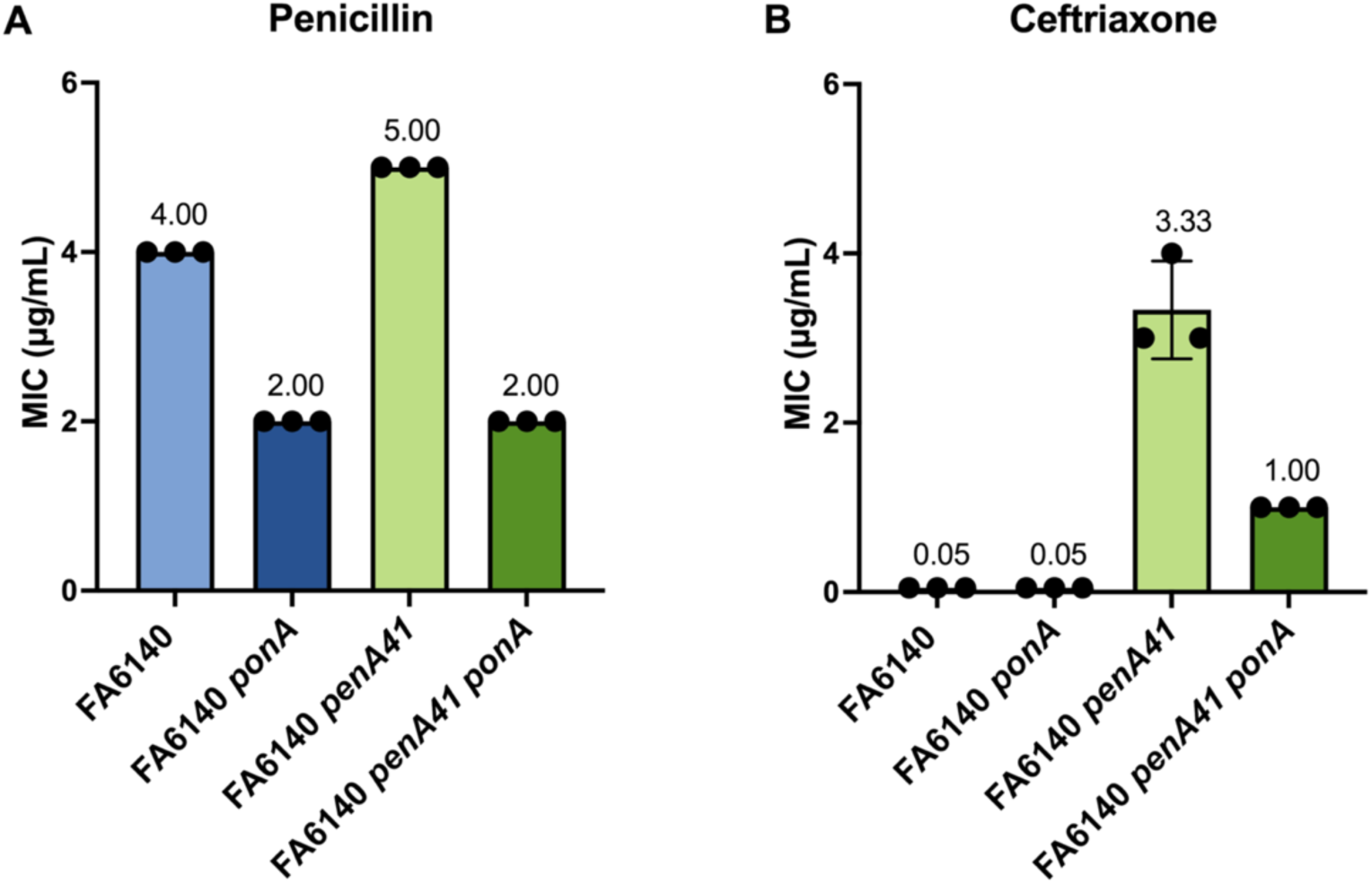
MICs of penicillin G and ceftriaxone for FA6140 transformants. The MICs reported above each bar are the median values (± the standard deviation) determined from three independent experiments. Data points within each bar represent the MIC values measured during each experiment.

## Discussion

In this study, we sought to understand the origins and role of the PBP1-L421P mutation (encoded by the *ponA*^L421P^ allele) in β-lactam antibiotic resistance in *N. gonorrhoeae.* Our data revealed that *ponA*^L421P^ is found exclusively in *N. gonorrhoeae* and its emergence in gonococcal isolates is likely due to it increasing the MIC_PEN_ above the clinical breakpoint of 2 µg/ml in CMRNG. While *ponA*^L421P^ has a modest fitness cost *in vivo* (29), there is no apparent fitness cost during growth *in vitro*. Our data suggest that this cost turns into a fitness benefit during growth in sub-MIC_PEN_ concentrations. Moreover, while penicillin G is no longer used to treat gonococcal infections, *ponA*^L421P^ also increases the MIC of ceftriaxone in strains harboring mosaic *penA41* alleles that confer ceftriaxone resistance. Thus, the increases in MIC_PEN_ (in FA6140) and MIC_CEF_ (in FA6140 *penA41*) that result from harboring *ponA*^L421P^ appear to have provided enough of an evolutionary advantage to remain in the population despite its modest fitness cost.

Mosaic *penA* alleles are defined by a sub-set of mutations that surround the active site of PBP2 and decrease the conformational dynamics of the PBP, which reduces acylation rates and increases MICs for various β-lactam antibiotics (15, 30, 31). These mutations emerged in *N. gonorrhoeae* through interspecies recombination events with *Neisseria* commensals. In contrast, *ponA*^L421P^ appears to have arisen spontaneously in *N. gonorrhoeae,* and, unlike resistance-conferring mutations in PBP2, L421P is located on a hinge that links the oligosaccharide-binding domain to the start of the transpeptidase domain in PBP1 and is over 20 Å from the active-site serine residue (Fig. 5). How the L421P mutation decreases acylation is unknown, and compared to mosaic PBP2 variants, the effects on *k_2_*/K_S_ are modest (8). The small decrease in acylation, however, could explain the 2-fold increase in the MIC_PEN_.

While *ponA*^L421P^ was the only mutation in FA6140 to increase the MIC_PEN_ to 4 μg/ml, another mutation, *ponA*^L421S^, increased the MIC_PEN_ to 2.5 μg/mL. However, to date *ponA*^L421S^ has not been documented in clinical isolates. This could be due to three reasons: 1) the L421P mutation requires only a single base change in the Leu-421 codon in *ponA* (C<u>T</u>G ➜ C<u>C</u>G), whereas the L421S mutation would require at least two base changes (CTG ➜ <u>TCT</u>/<u>TCC</u>/<u>TCA</u>/<u>TC</u>G/<u>AGT</u>/<u>AGC</u>); 2) strain FA6140 *ponA*^L421S^ is less fit *in vitro* compared to other strains; and 3) the increase in resistance is insufficient to overcome its fitness cost.

Following the six serial passages of the 16 mutants, we sequenced the genomes of the 36-hour cultures from all three biological replicates to be sure that the results were not skewed by the acquisition of random mutations, particularly under antibiotic selective pressure. No consistent or biologically relevant mutations were detected, with the exception of a single frameshift mutation in *pilQ* observed in the culture supplemented with 1.0 μg/mL of penicillin during trial 1. PilQ increases the MICs of a range of antimicrobials in *N. gonorrhoeae* strains that also have the *mtr* and *porB* resistance determinants (18, 19), which decrease the concentration of antibiotics in the periplasm by increasing efflux and decreasing influx of antibiotics, respectively. This implies that permeation through the PilQ secretin is low and does not impact the MIC unless both *mtr* and *porB* determinants are present, as they are in FA6140 (3, 8, 18, 22, 35). Based on the increase in read counts for *ponA*^L421S^ at 24 hours, we surmise that the PilQ mutation arose in FA6140 *ponA*^L421S^ sometime between 18-24 hours of growth, which increased its MIC_PEN_ above that of FA6140 *ponA*^L421P^ and allowed it to survive in higher penicillin concentrations. However, this is an outcome that would likely occur only during *in vitro* growth, as expression of type IV pili (of which PilQ is an essential component) is important for *N. gonorrhoeae* survival during pathogenesis (36). This is supported by the absence of inactive PilQ mutants identified in clinical isolates (37, 38).

We recently reported that *N. gonorrhoeae* lineages harboring *ponA*^L421P^ between 1993 and 2013 were associated with a modest fitness disadvantage, compared to the baseline type (lineages that do not carry any resistance determinants), suggesting that *ponA*^L421P^ imposes a fitness cost (29). These data were consistent with results from competitive co-infections between *ponA*– and *ponA*^L421P^-containing strains of FA19 and FA6140 (29). Despite this observed fitness cost, we showed here that *ponA*^L421P^ remains prevalent in the gonococcal population and is present in over 90% of isolates that also harbor a mosaic *penA* allele (as classified by the NG-STAR database). Why is *ponA*^L421P^ found so frequently in strains with a mosaic *penA* allele? There are several possible explanations for this: 1) mosaic *penA* alleles emerged by transformation almost exclusively into penicillin-resistant strains that already had *ponA*^L421P^, as these strains also contain other resistance determinants and presumably compensatory mutations that lessen their fitness cost; 2) with penicillin resistance at or near 100% in Asia (39), where ceftriaxone resistance has been emerging, the link between *ponA*^L421P^ and ceftriaxone resistance may reflect the gonococcal diversity/population structure in the places experiencing the most selection for ceftriaxone resistance; and 3) *ponA*^L421P^ confers additional resistance to ceftriaxone and thus provides a benefit to these strains, given the widespread use of ceftriaxone to treat current infections. We tested the latter possibility by determining the MIC_CEF_ values of FA6140 and FA6140 *ponA* also harboring the mosaic *penA41* allele from H041, which confers ceftriaxone resistance to these strains. Importantly, the MIC_CEF_ drops ∼3-fold when *ponA*^L421P^ is replaced with *ponA*, indicating that *ponA*^L421P^ confers additional resistance to ceftriaxone and potentially explaining, at least in part, why *ponA*^L421P^ is so prevalent in strains also harboring mosaic *penA* alleles, particularly with mosaic *penA* alleles that confer the largest increases in MIC_CEF_.

The MIC data we present in this study can be best interpreted by knowing which PBP is being targeted by penicillin and ceftriaxone (Fig. 9). The acylation constant *k_2_*/K_S_ for each antibiotic determines which PBP first becomes acylated as the antibiotic concentration increases (the higher the *k_2_*/K_S_, the lower the concentration of antibiotic needed to inhibit the PBP). Thus, *k_2_*/K_S_ and the MIC have an inverse relationship, and the essential PBP that first becomes acylated as the antibiotic concentration increases is the killing target for that antibiotic and determines the MIC. For penicillin, the killing target for FA6140 is likely a combination of PBP2^FA6140^ and PBP1^L421P^, but when PBP1^L421P^ is replaced by PBP1, the MIC decreases because PBP1 becomes the exclusive killing target of penicillin. We showed previously that in 35/02, a ceftriaxone-intermediate resistant (and penicillin-resistant) gonococcal isolate, reversion of *ponA*^L421P^ to *ponA* did not alter MIC_CEF_ values (40). In contrast, expression of PBP2^H041^ in FA6140, which has a much lower *k_2_*/K_s_ for ceftriaxone than PBP2^35/02^, shifts the killing target to PBP1^L421P^. Thus, exchanging PBP1^L421P^ for PBP1 in the FA6140 genetic background decreases MIC_CEF_ by 3-fold, suggesting that when strains acquire a “strong” mosaic PBP2 variant, e.g. PBP2^H041^, that has a sufficiently low acylation rate constant for ceftriaxone, the target shifts from PBP2 to PBP1 and *ponA*^L421P^ confers a selective advantage over *ponA*.

**FIG 9.**
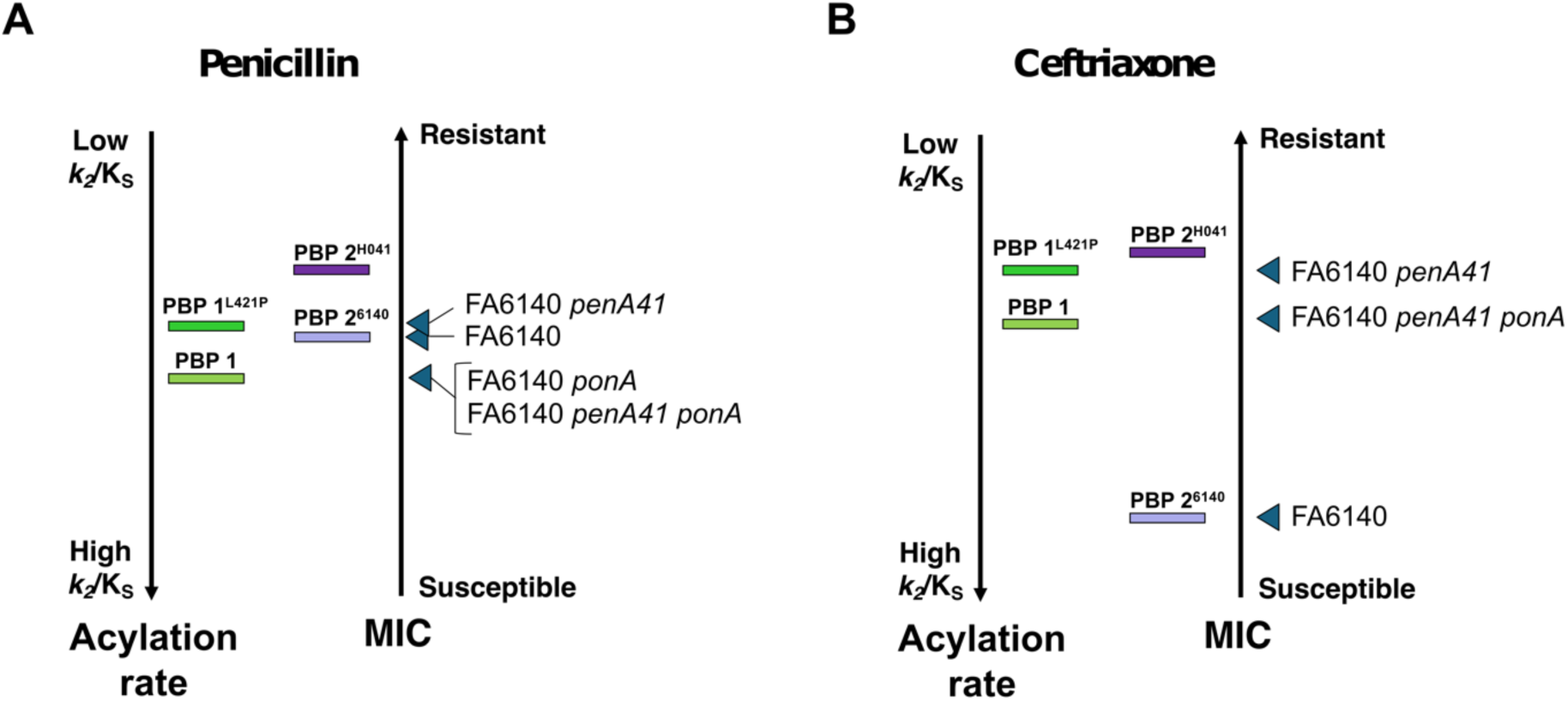
Diagram outlining the drug targets for penicillin and ceftriaxone in gonococcal strains with different *penA* and *ponA* alleles. PBP1 variants are marked by green rectangles and PBP2 variants are marked as purple rectangles. The left axis indicates the acylation rate, and the right axis indicates the MIC of the indicated strain. Arrows point to the killing target for each of the indicated strains. A). Penicillin has a low *k_2_*/K_S_ for all PBP2 variants, so it kills gonococcal cells by inhibiting PBP1. Replacing PBP1^L421P^ with PBP1^WT^ decreases the MIC. B). Ceftriaxone normally kills by inhibiting PBP2, but PBP2^H041^ has a much lower *k_2_*/ K_S_ and this shifts the killing target to PBP1 (and maybe some PBP2^H041^). Replacing PBP1^L421P^ with PBP1^WT^ decreases the MIC.

In conclusion, this is the first study to investigate the origins of *ponA*^L421P^ in the context of penicillin resistance and address why it is maintained in the *N. gonorrhoeae* population. During the 40+ years of penicillin use, *N. gonorrhoeae* strains acquired β-lactamases, as well as mutations in PBP1, PBP2, *mtr*, and *porB*, which eventually increased the MIC_PEN_ enough to result in its removal as a treatment option. When expanded-spectrum cephalosporins (ESCs) were introduced to treat gonococcal infections, mosaic PBP2 variants emerged (because PBP2 was the exclusive killing target of ESCs), in existing penicillin-resistant strains, resulting in the proliferation of mosaic *penA* alleles that increased resistance to ESCs. Over time, the MICs of ESCs for gonococcal isolates increased as the mosaic *penA* alleles picked up additional mutations, and the killing target shifted to PBP1, allowing the L421P mutation to further increase the MIC. These findings reinforce the need to identify antibiotics that are PBP1-selective and thus can avoid the highly mutated mosaic PBP2 variants that are compromising the use of ESCs. β-lactam antibiotics that target both PBPs or a combination of two β-lactam antibiotics that target PBP2 and PBP1 may provide a new strategy for treating strains with mosaic *penA* alleles.

## Methods

### Strains and plasmids

Strains, plasmids, and primers used in this study are listed in Tables S1 and S2. To aid in subsequent screening for generating random mutations at codon-421, we transformed FA6140 with the *ponA* allele from FA19, which naturally has a PstI site encompassing codons 421 and 422. A portion of the *ponA* gene (bp 956 – 2400) from FA19 genomic DNA was amplified via PCR and transformed into FA6140 via spot transformation, performed as described (41). Colonies were screened for *ponA* by PCR amplification of *ponA* and digestion with PstI. To generate FA6140 and FA6140 *ponA* harboring the mosaic *penA* allele (*penA41*) from the ceftriaxone-resistant strain H041, piliated cells were transformed with pUC18us-*penA41* plasmid containing *penA41* starting at bp 133 and ending 200 bp downstream *penA41* to facilitate recombination. Transformants that acquired *penA41* were selected on GCB agar plates containing 1.0 !g/mL cefixime. All transformants were verified by PCR and Sanger sequencing.

Descriptions of the cloning plasmids used to create the library of isogenic FA6140 *ponA*^L421X^ mutants can be found in Table S3. pUC18us-*ponA*^421X^-Ω contains a portion of the *ponA* gene (from bp 831 to 54 bp downstream of the stop codon) harboring randomized codons to introduce amino acid substitutions at codon-421, the *aad1* resistance cassette (Ω) conferring spectinomycin/streptomycin resistance (42), and 531 bp of sequence downstream of *ponA* to facilitate recombination. We employed overlap-extension (43) PCR mutagenesis to introduce amino acid substitutions at codon-421 in *ponA* fragments, using the primers listed in Table S2. pUC18us-*ponA*^421X^-Ω plasmids were created using the NEBuilder HiFi DNA Assembly Master Mix and transformed into *Escherichia coli* XL-10 Gold cells, according to manufacturer’s protocol. Clones were screened by PCR amplification of the *ponA* gene followed by Sanger sequencing. Plasmids containing unique mutations were isolated using a QIAPrep miniprep kit (Qiagen). Plasmids (Table S3) were transformed into FA6140 *ponA* by allelic exchange, and transformants were selected on GCB agar plates containing 25 !g/mL spectinomycin. Individual colonies from each transformation were screened by PCR by amplifying the *ponA* gene and digesting with PstI (loss of PstI indicated the presence of the mutation). All clones were verified by Sanger sequencing.

PCR reaction conditions were as follows; 94°C for 30s (denaturation), 60°C for 30s (annealing), and 72°C for 60s (elongation) for 35 cycles, and fragments were purified using the QIAquick PCR Purification kit (Qiagen) as per manufacturers guidelines. All gonococcal strains were propagated on solid GCB agar containing Kellogg’s supplement I (38) and 12 mM Fe(NO_3_)_3_ for 18 to 20 h at 37°C in a 5% CO_2_-enriched humidified atmosphere. Serial passaging was minimized to reduce the risk of acquiring secondary mutations.

### Bacterial growth

FA6140 *ponA*^L421X^ mutants were grown in liquid culture to verify individual growth phenotypes (Table S4). Nonpiliated bacterial colonies were swabbed from GCB agar grown for 16 h and inoculated into GCB (per liter: 15 g protease peptone 3, 4 g K_2_HPO_4_, dibasic, 1 g KH_2_PO_4_, 5 g NaCl) supplemented with Kellogg’s supplement I (38) and 12 mM Fe(NO_3_)_3_ at a starting optical density at 600 nm (OD_600_) of 0.08. Cultures were shaken at 180 rpm at 37°C in a 5% CO_2_-enriched atmosphere, and bacterial growth was assessed by measuring the OD_600_ at hourly intervals for a total of 8 h. Experiments were repeated in biological triplicate, and statistical significance between individual strains was determined by using a repeated-measures 2-way analysis of variance (ANOVA) with Tukey’s multiple comparisons. Differences in growth rate were measured by comparing the average time to reach an OD_600_ of 0.8 for each strain as previously described (44). Results were compared using a one-way analysis of variance to determine overall significance followed by Dunnett’s multiple comparison test to determine significance between individual strains.

### *Neisseria* sequence diversity analysis

The datasets used to identify polymorphisms and assess overall sequence diversity of the *ponA* gene across all *Neisseria* species were all exported from the *Neisseria* page of the PubMLST data base (last accessed September 30, 2024) (45). Allele mining analysis of the *ponA* gene was performed as described (46). An individual search of the profiles was made via the locus/sequence definitions tool to identify the locus assigned to *ponA* (NEIS0414). Data was exported from the two-field breakdown section of the *Neisseria* isolate database as a Microsoft Excel file, which provided a list of unique *ponA* allelic profiles and a count for how many isolates possess that unique allele within each *Neisseria* spp. Microsoft Excel was used to calculate the number of isolates that possess each allele. Mutations identified in the *ponA* alleles from *N. gonorrhoeae* isolates with frequency greater than 5% were mapped onto the predicted PBP1 structure, which was generated using Alphafold (47). The PDB file for PBP1 from *N. gonorrhoeae* (UniProt: O05131) was downloaded from the AlphaFold Protein Structure Database (AFDB) (48, 49) and mutations detected frequencies greater than 5% were mapped to the structure using Pymol (2.0.7) (50). *ponA* sequences from PubMLST were exported using the script available in the allele/sequence definition database. Molecular Evolutionary Genetics Analysis software, version 7 (MEGA7) (51) was used to translate the sequences and perform a multiple sequence comparison by log-expectation (MUSCLE) (52). Sequence logo images were created using WebLogo (53). MUSCLE alignments of unique *ponA* alleles from PubMLST were uploaded to WebLogo, background composition of proteome was set to equiprobable, and y-axis units were set to bits.

### Genomic Analyses

*Genome assembly and typing.* Publicly available *N. gonorrhoeae* genomic data with available MICs of penicillin were downloaded from the European Nucleotide Archive (54–72). Resistance-associated SNPs were determined from variant calls produced by Pilon v 1.23 (73) after mapping to the NCCP11945 reference genome (NC_011035.1) using BWA-MEM v 0.7.17 (74). Mosaic alleles and the presence or absence of resistance-associated accessory genes (e.g. *bla*_TEM_) were identified using blastn v 2.14.0 (75) searches of *de novo* assemblies generated with SPAdes v 3.12.0 (76). Mosaic *penA* alleles were typed according to the NG-STAR database (77).

Genomic data was included if the total *de novo* assembly length was within the range of expected lengths for *N. gonorrhoeae* (1.89 Mb – 2.31 Mb) and that the number of contigs was less than 250. We also required more than 40X coverage of the reference genome, that at least 80% of reads mapped to the reference genome, and less than 12% of positions in the reference genome had ambiguous calls defined as at least 90% of reads supporting either the reference or alternate nucleotide. Lastly, we removed isolates with ambiguous calls in genes that contribute to penicillin resistance, including *ponA, penA*, and the *mtr* operon.

*Phylogenetic analysis.* An initial alignment and core genome phylogeny was estimated using ParSNP v 2.0.5 (78) from 10,282 gonococcal *de novo* assemblies. To reduce the computational resources needed for a recombination-corrected phylogenetic analysis, we used Treemmer v 0.3 (79) to prune the ParSNP tree to 99.9% of the original tree length, removing the most closely related pairs of isolates from our dataset. For this representative subset of 6,082 isolates, pseudogenomes were generated from variant calls and concatenated to produce an alignment for recombination detection and phylogenetic analysis with Gubbins v 3.3.3 (80). Phylogenetic trees were visualized using iTOL (81). We additionally pruned the dataset to remove isolates encoding *blaTEM*, leaving a representative dataset of 5,192 isolates to investigate the distribution of *ponA* among CMRNG.

### Antimicrobial susceptibility testing

All MIC values were determined by the agar dilution method exactly as described previously (34). Briefly, FA6140 transformants were grown overnight on GCB plates, resuspended in GCB broth at a final concentration of ∼1 × 10^7^ colony forming units (CFU)/ml. Aliquots (5 μl) of each strain were spotted onto GCB plates containing antibiotics at <2-fold dilutions. The MIC was defined as the lowest concentration of antibiotic on which fewer than 5 colonies appeared after 24 hrs of incubation.

### *In vitro* serial passaging

Non-piliated colonies from each individual FA6140 *ponA*^L421X^ strain (Table 1) were harvested and used to prepare a starting inoculum, which consisted of an equal ratio (verified through Illumina sequencing) of all 16 strains. An equal volume of the starting inoculum was used to inoculate liquid cultures at 0 h at an OD_600_ of 0.08. Three cultures were supplemented with a different sub-MIC concentration of penicillin (0.25, 0.5, and 1.0 μg/mL), and one culture remained penicillin-free. Penicillin was added at hour 0 at the specified concentration and was maintained throughout each of the six 6-hr growth periods. After inoculation, cultures grew for 6 hours, at which time the OD_600_ of each culture was measured to calculate the volume of cells needed to inoculate the next culture at an OD_600_ of 0.08, and an aliquot was removed for analysis. This passaging procedure was performed every 6 h for a total of 36 h. At every 6-hour timepoint, the cultures were checked for contamination by dipping a sterile swab in the media and streaking it out on a GCB agar plate. This experiment was repeated 3 separate times. All liquid cultures consisted of 40 mL of GCB + supplements and were shaken at 180 rpm at 37°C in a 5% CO_2_-enriched atmosphere.

### Genomic DNA extraction

DNA was extracted from cells collected every 6 h during the *in vitro* serial passaging experiments using the Promega Wizard genomic DNAs purification kit (Promega, Madison, WI) according to the manufacturer’s protocol. Extracted gDNA is from a mixture of cells that represent the ratio of all sixteen FA6140 *ponA*^L421X^ mutants growing in the culture at that time point. DNA concentrations were measured on the Invitrogen Qubit 2.0 Fluorometer. Samples were prepped using the Qubit™ dsDNA Quantification Assay Kits according to the manufacturer’ guidelines (Invitrogen).

### PCR and DNA sequencing of FA6140 *ponA*^L421X^ mutants during *in vitro* serial passaging

*Library preparation.* Fragments for the library were prepped in a series of two PCR reactions. The first PCR reaction (PCR 1; 20 cycles of 94°C for 30 s, 60°C for 30 s, and 72°C for 15 s) used genomic DNA extracted from cells collected during the serial *in vitro* passaging experiments as a template. Primers were designed to target a region of the *ponA* gene (bp 1212-1309), which encompasses codon-421, and to attach the appropriate adaptor sequences to the 5’ and 3’ ends of each amplified fragment. The amplicons obtained were then purified and used as templates in the second PCR (PCR 2; 12 cycles of 94°C for 30 s, 60°C for 30 s, and 72°C for 15 s). Primers used in PCR 2 target the 5’ and 3’ Illumina adaptor sequences from PCR 1. Barcoding was achieved by adding a unique, 5 bp sequence to the 5’ end of fragments to signify the time point it refers to. A unique index sequence was attached to the 3’ ends of the fragments to mark the penicillin concentration of the culture from which the fragment was obtained. All PCR fragments were extracted from 3% agarose gels and purified using the QIAquick Gel Extraction Kit (QIAGEN) according to manufacturers’ guidelines. Concentrations were measured on the Invitrogen Qubit 2.0 Fluorometer.

*Illumina sequencing.* Purified fragments from PCR 2 were pooled and sequencing was performed on an Illumina Next-seq 1000. This was done three separate times, once per *in vitro* serial passaging trial.

*Data analysis.* Read files generated from our libraries contained between 10 and 70 million reads. The reads in this library are essentially all genetically identical copies of the amplified region of *ponA*, except for codon-421, meaning each read correlates to a strain from the FA6140 *ponA*^L421X^ library, based on the amino acid at codon-421. Data were parsed using a UNIX script to count codons and condition specific barcodes. This process yielded a table for each timepoint, listing all of the codons identified at position-421, and a count of how many reads have that codon. Read counts across time points were normalized to the total number of reads in the file. Only informative reads, containing the expected codons at the expected positions, were analyzed to limit normalization and counting based on uncertain read calls. After filtering, we retained ∼99% of reads that were informative. Bar graphs were generated by normalizing codon counts at 36 hr to the strain’s inoculum value (codon count at 0 h). Since this is a zero-sum competitive experiment, any gain in codon counts by one isolate directly corresponds to a loss in codon counts by the others.

### Whole genome sequencing and assembly

Whole genome sequencing was performed by SeqCenter (Pittsburgh, PA) on genomic DNA from the inoculum and the 36 h timepoint. Illumina sequencing libraries were prepared using the tagmentation-based and PCR-based Illumina DNA Prep kit and custom IDT 10 bp unique dual indices (UDI) with a target insert size of 280 bp. No additional DNA fragmentation or size selection steps were performed. Illumina sequencing was performed on an Illumina NovaSeq X Plus sequencer in one or more multiplexed shared-flow-cell runs, producing 2 x 151 bp paired-end reads. Demultiplexing, quality control and adapter trimming was performed with bcl-convert1 (v4.2.4). Variant calling analysis on the reads was also performed by SeqCenter, using their standard methods. Illumina-generated 2 x 151 bp paired-end read data was used as the input for variant calling against the reference genome of gonococcal clinical isolate, FA6140 (82), the parental strain used to create the library of mutants (GCF_001047255.1). Variant calling was carried out by SeqCenter using BreSeq1 under default settings.

## Data availability

This publication made use of the PubMLST website (http://pubmlst.org/) developed by Keith Jolley (45) and sited at the University of Oxford. The development of that website was funded by the Welcome Trust. Data was retrieved and is available from https://pubmlst.org/organisms/neisseria-spp. Data supporting the findings are included in the manuscript. All metadata and quality metrics for isolates included in the datasets for our genomic and phylogenetic analysis are available at https://github.com/mortimer-lab/ponA421. Experimental data, raw FASTQ files, and tables with codons and counts for each Illumina sequencing trial, as well as the code used to generate these tables are available upon request.

## Supplemental Material

TABLE S1

TABLE S2

TABLE S3

TABLE S5

FIG S1

FIG S2

FIG S3

FIG S4

FIG S5

## Acknowledgements

The contents of this article are solely the responsibility of the authors and do not necessarily represent the official views of the Uniformed Services University of the Health Sciences, the Department of Defense, the University of North Carolina, Harvard University, and the University of Georgia. This work was supported by grants AI153521 (YHG, AEJ, and RAN) and AI164794 (RAN), as well as the Pharmacological sciences T32 Training Program award, GM135095 (GG), and the National Institute of Allergy and Infectious Disease (NIAID) STI/HIV T32 Training Program award, T32AI007001 (GG). The funders had no role in study design, data collection and interpretation, or the decision to submit the work for publication.

GG and RAN designed and conceptualized the research, GG performed all experiments, analyzed data, provided opinions, and wrote the manuscript. TDM performed genome assembly and typing and phylogenetic analysis, reviewed and edited the article, and provided opinions. BG and DD assisted with Illumina sequencing, providing insightful advice on experimental design and data analysis. ALV, AEJ, and YHG provided input and opinions during the course of this study, and critical review of the manuscript. Supervision, funding acquisition, and article revisions provided by RAN.

