## Supplemental Material for "The role of the L421P mutation in Penicillin-Binding Protein 1 (PBP1) in the evolution of chromosomally mediated penicillin resistance in *Neisseria gonorrhoeae*"

**Table S1.** Strains and plasmids used in this study.

| Strain / Plasmid Name | Description[a] | Reference |
|---|---|---|
| **FA19** | Wild type, antibiotic-susceptible *N. gonorrhoeae* lab strain. | (1) |
| **FA6140** | Penicillin-resistant *N. gonorrhoeae* clinical isolate that naturally harbors the *ponA1* allele. | (2) |
| **H041** | Multi-drug-resistant *N. gonorrhoeae* clinical isolate. Displays high-level resistance to ceftriaxone and penicillin. | (3) |
| **FA6140 *ponA*** | FA6140 x *ponA* (FA19) PCR; replaces *ponA*$^{L421P}$ with *ponA* | This study |
| **FA6140 *penA41*** | FA6140 x pUC18us-*penA41*; replaces *penA6140* with *penA41* | This study |
| **FA6140 *penA41 ponA*** | FA6140 *ponA* x pUC18us-*penA41*; replaces *penA6140* with *penA41* | This study |
| **pUC18us-*penA41***[b] | Plasmid pUC18us with full-length *penA41* from gonococcal isolate H041 | This study |

[a] Transformants are shown as recipient strain x donor DNA, produced as described in Materials and Methods. PCR signifies a PCR product was used to transform.
[b] pUC18us contains the 10 bp gonococcal uptake sequence preceding the multiple cloning site.

**Table S2** Oligonucleotides and plasmids used for cloning in this study.

| Primer name | Sequence (Annealing) | Description |
|---|---|---|
| 5'pUC18US_*ponA*_HiFi | TGCCAAGCTGGCCGTCTGAAAAGCTTATATGGTGAAGATGCCTATACG | Forward primer to amplify the *ponA* allele from Ng DNA for ligation into pUC18(Ω) |
| 3'pUC18US_ponA_HiFi | AGCGTGCATAATAAGCCCTATCTAGAGCCAAATCTAAAATGCCGTC | Reverse primer to amplify the *ponA* allele from Ng DNA for ligation into pUC18(Ω) |
| 5'*ponA*956 | GCGGTGCGGAAAACTATATC | Forward primer to Sanger sequence the *ponA* allele of transformed *Ng* |
| 3'*ponA* | TGTTTGACAGCTTATCATCGATAAGCTTTTAAAACAGGGAATCCAACTGC | Reverse primer to Sanger sequence the *ponA* allele of transformed *Ng* |
| 5'*ponA*421random | CAGGGGGCTTTGGTTTCGCTGGA | Forward primer to amplify the *ponA* allele from *Ng* DNA, primer anneals to *ponA* at bp 1264-1286. |
| 3'*ponA*421random | TCCAGCGAAACCAAAGCCCCCTGNNNCAACGGCTCTTGAACCACCG | Reverse primer to amplify the *ponA* allele from *Ng* DNA and introduce randomized codon at position 421, primer anneals to *ponA* at bp 1241-1286. |
| 5'penA45-pUC18 | AAGCTGGCCGTCTGAAAAGCTTCGCGGGCTGTATCTGCAGACGGTAA | Forward primer to amplify the *penA41* allele from *Ng* DNA for ligation into pUC18 |
| 3'murE19-87-pUC18 | CAGCTATGACCATGATTACGAATTCCGGCACGGTGTTTCAAATCT | Reverse primer to amplify the *penA41* allele from *Ng* DNA for ligation into pUC18 |
| 5'penA-45aa | CGCGGGCTGTATCTGCAG | Forward primer to Sanger sequence the *penA41* allele, starting at position 45, from pUC18us-*penA41*-transformed *Ng* |
| 5'H041-317aa | ATTGCCAAAGCGCTGGATTC | Forward primer to Sanger sequence the *penA41* allele, starting at position 317, from pUC18us-*penA41*-transformed *Ng* |
| 5'pUC18_40 | GCCAGGGTTTTCCCAGTCACGA | Forward primer to Sanger sequence pUC18us-Ω plasmid constructs |
| 3'pUC18_Rev | GAGCGGATAACAATTTCACACAGG | Reverse primer to Sanger sequence pUC18us-Ω plasmid constructs |

**Table S3** Cloning plasmids used to create each FA6140 *ponA*$^{L421X}$ Mutant.

| Transformation Plasmid[a] | Amino Acid at PBP1$^{421}$ | Codon at PBP1$^{421}$ | Reference |
|---|---|---|---|
| **pUC18us-** ***ponA*$^{421X}$-Ω**[b] | - | NNN | This study |
| **pUC18us-** ***ponA*$^{421L}$-Ω** | Leucine (L) | CTG | This study |
| **pUC18us-** ***ponA*$^{421P}$-Ω** | Proline (P) | CCG | This study |
| **pUC18us-** ***ponA*$^{421A}$-Ω** | Alanine (A) | GCC | This study |
| **pUC18us-** ***ponA*$^{421R}$-Ω** | Arginine (R) | AGA | This study |
| **pUC18us-** ***ponA*$^{421C}$-Ω** | Cysteine (C) | TGC | This study |
| **pUC18us-** ***ponA*$^{421N}$-Ω** | Asparagine (N) | AAC | This study |
| **pUC18us-** ***ponA*$^{421E}$-Ω** | Glutamic Acid (E) | GAA | This study |
| **pUC18us-** ***ponA*$^{421G}$-Ω** | Glycine (G) | GGT | This study |
| **pUC18us-** ***ponA*$^{421H}$-Ω** | Histidine (H) | CAT | This study |
| **pUC18us-** ***ponA*$^{421Q}$-Ω** | Glutamine (Q) | CAG | This study |
| **pUC18us-** ***ponA*$^{421W}$-Ω** | Tryptophan (W) | TGG | This study |
| **pUC18us-** ***ponA*$^{421Y}$-Ω** | Tyrosine (Y) | TAT | This study |
| **pUC18us-** ***ponA*$^{421S}$-Ω** | Serine (S) | AGT | This study |
| **pUC18us-** ***ponA*$^{421F}$-Ω** | Phenylalanine (F) | TTC | This study |
| **pUC18us-** ***ponA*$^{421I}$-Ω** | Isoleucine (I) | ATC | This study |
| **pUC18us-** ***ponA*$^{421T}$-Ω** | Threonine (T) | ACT | This study |

[a]All transformation plasmids are pUC18-us containing bp 831-2400 of *ponA*, 54 bp downstream of *ponA*, the *aad1* resistance cassette (Ω), and 531 bp of additional downstream sequence to aid in transformation. Each plasmid contains a different codon at position 421 and was cloned into the parental strain FA6140 *ponA* (FA6140 with a wt *ponA* allele).

[b] Plasmid used to generate the rest of the plasmids in this table, as described in Materials and Methods.

**Table S4** *In vitro* growth kinetics of the FA6140 *ponA*$^{L421X}$ mutants, compared to the parent strain, FA6140.

| Strain Name[a] | Amino Acid at PBP1$^{421}$ | Time (min) to reach OD600 of 0.8 ± SE | P-value for Strain vs FA6140[b] | Summary |
|---|---|---|---|---|
| **FA6140** | P | 294.1 ± 8.4 | - | - |
| FA6140 *ponA*$^{L421P}$ | P | 285.0 ± 27.1 | 0.9993 | ns |
| FA6140 *ponA*$^{L421L}$ | L | 269.9 ± 47.7 | 0.9842 | ns |
| FA6140 *ponA*$^{L421A}$ | A | 248.3 ± 44.1 | 0.7203 | ns |
| FA6140 *ponA*$^{L421R}$ | R | 236.1 ± 32.2 | 0.4935 | ns |
| FA6140 *ponA*$^{L421C}$ | C | 272.5 ± 4.9 | 0.9902 | ns |
| FA6140 *ponA*$^{L421N}$ | N | 252.7 ± 24.4 | 0.7985 | ns |
| FA6140 *ponA*$^{L421E}$ | E | 229.8 ± 22.6 | 0.3899 | ns |
| FA6140 *ponA*$^{L421G}$ | G | 218.9 ± 18.9 | 0.2457 | ns |
| FA6140 *ponA*$^{L421H}$ | H | 253.1 ± 32.4 | 0.8055 | ns |
| FA6140 *ponA*$^{L421Q}$ | Q | 236.6 ± 24.0 | 0.5029 | ns |
| FA6140 *ponA*$^{L421W}$ | W | 268.0 ± 20.8 | 0.9749 | ns |
| FA6140 *ponA*$^{L421Y}$ | Y | 290.2 ± 15.1 | 0.9996 | ns |
| FA6140 *ponA*$^{L421S}$ | S | 272.8 ± 9.0 | 0.9904 | ns |
| FA6140 *ponA*$^{L421F}$ | F | 287.7 ± 8.4 | 0.9994 | ns |
| FA6140 *ponA*$^{L421I}$ | I | 254.1 ± 14.7 | 0.8222 | ns |
| FA6140 *ponA*$^{L421T}$ | T | 223.8 ± 29.7 | 0.3050 | ns |

[a] All strains are in the FA6140 background, and each contain a different amino acid at residue 421 of *ponA* (PBP1).

[b] Overall significance was determined by a one-way analysis of variance, followed by Dunnet's multiple comparisons test to determine significance between individual strains.

**Table S5** Percentage of *Neisseria* isolates that harbor leucine or proline at residue 421 in *ponA*.

| | ***Neisseria* Isolates Harboring[b]:** | |
|---|---|---|
| ***Neisseria Spp.*[a]** | **Leucine at residue 421 in *ponA*** | **Proline at residue 421 in *ponA* (*ponA$^{L421P}$*)** |
| *N. gonorrhoeae* (n = 20,513) | 10,351 (50.46%) | 10,162 (49.54%) |
| *N. meningitidis* (n = 75,612) | 75,612 (100%) | 0 (0%) |
| Commensal isolates[c] (n = 1,604) | 1,604 (100%) | 0 (0%) |

[a] Isolate collections are from PubMLST database (n = total number of isolates included in analysis).

[b] Number (%) of isolates that harbor specified amino acid at residue 421 of PBP1 (encoded by *ponA*).

[c] Commensal isolates include strains from; *N. bergeri, N. animalis, N. basseii, N. benedictiae, N. blantyrii, N. cinereal, N. lactamica, N. maigaei, N. mucosa, N. oralis, N. polysaccharea, N. subflava, N. uirgultaei, N. viridiae.*

**FIG S1** Sequence logo diagrams comparing the conservation of PBP1 from various *Neisseria* isolates. Diagram generated by WebLogo using MUSCLE aligned *ponA* sequences. Sequences were taken from PubMLST database and represent all the unique *ponA* alleles documented for each *Neisseria* species group, A). *N. gonorrhoeae* (n=172), B). *N. meningitidis* (n=897), and C). *Neisseria spp.* (n=369). The *x*-axis indicates the position of the amino acid and the *y*-axis shows the raw residue frequencies as bits. The overall height of the stack indicates the sequence conservation at that position. Columns with many gaps or unknown residues are narrow.

**A). *N. gonorrhoeae***

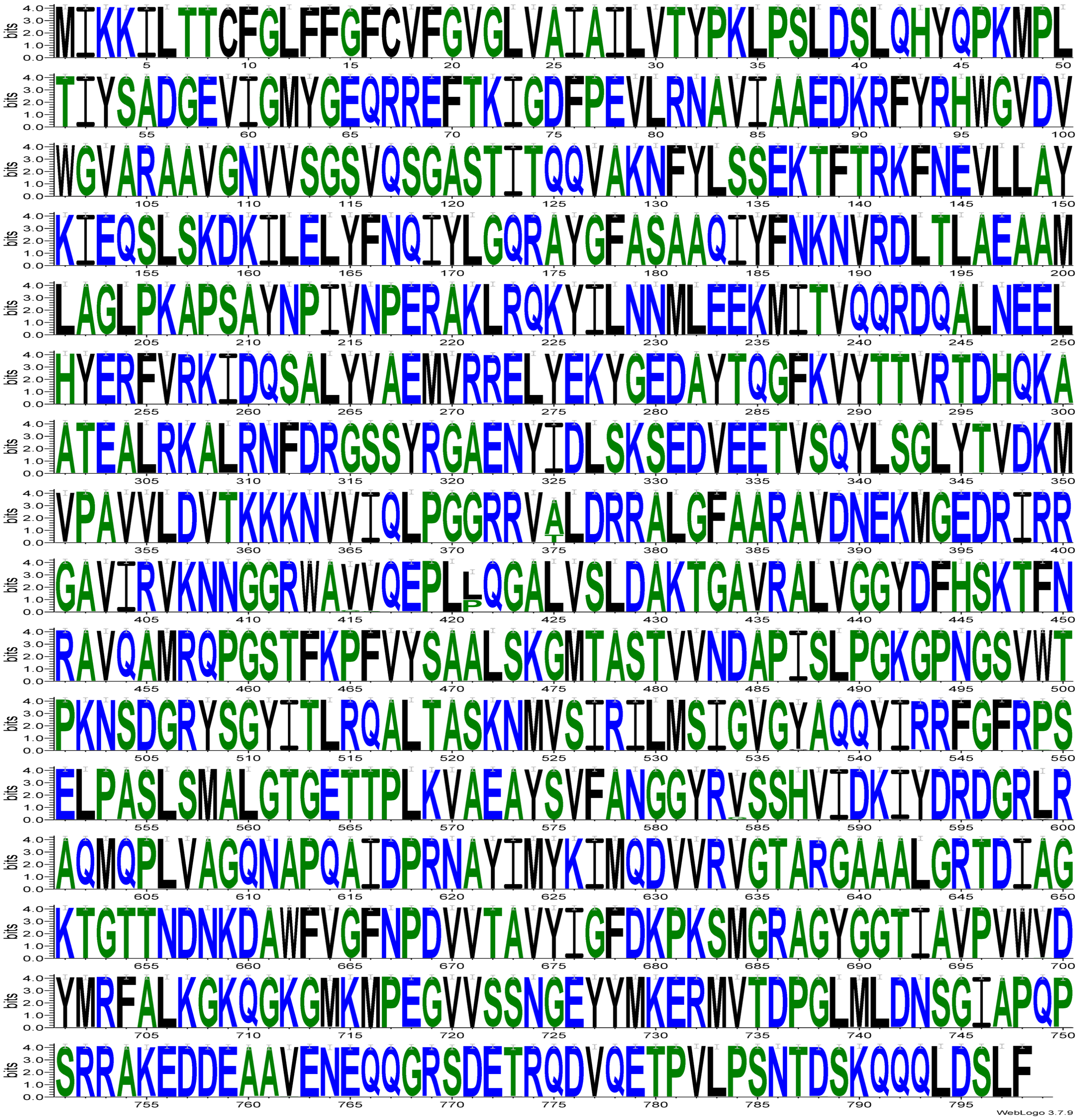

### B). *N. meningitidis*

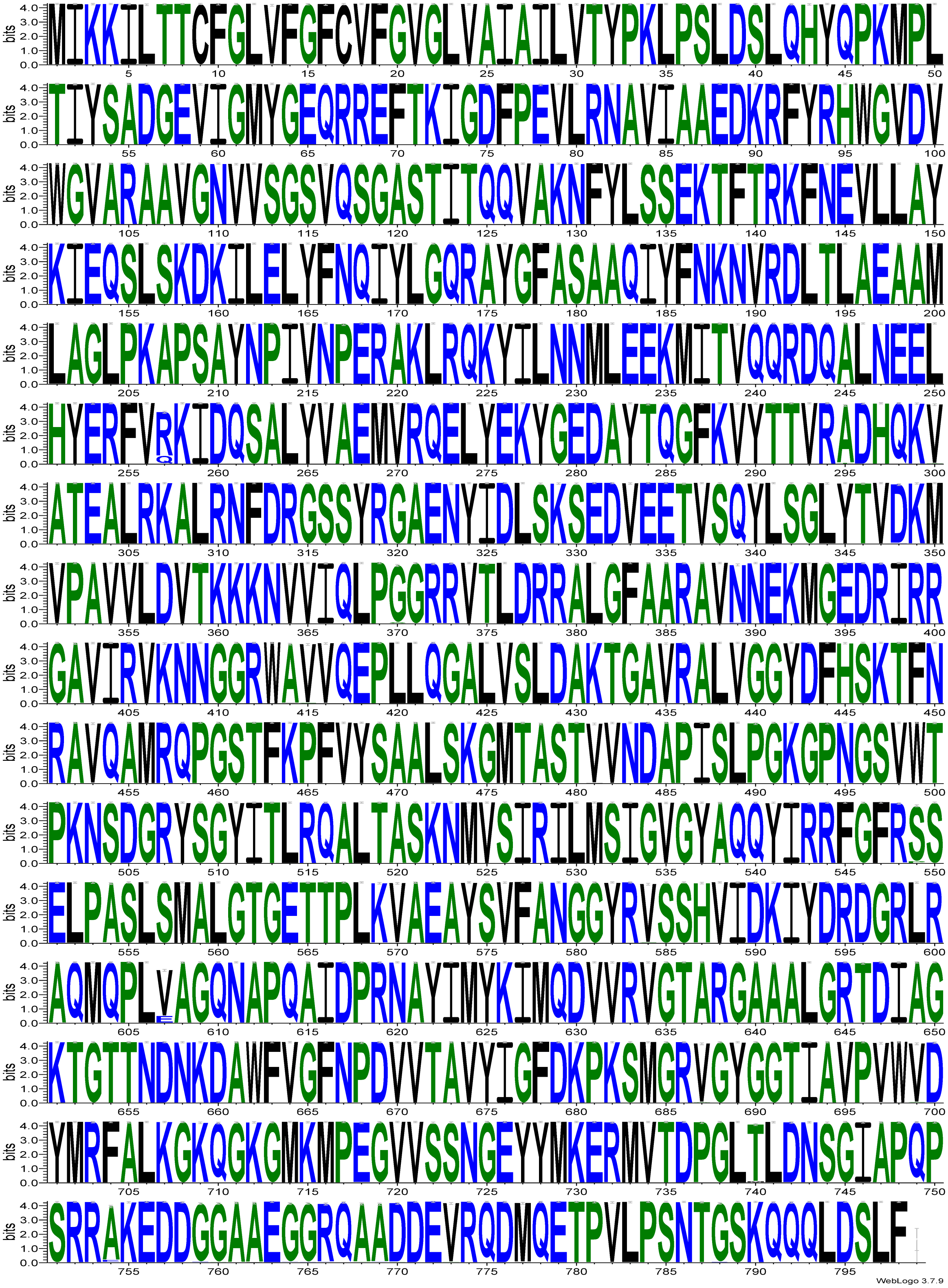

C). *Neisseria spp.*

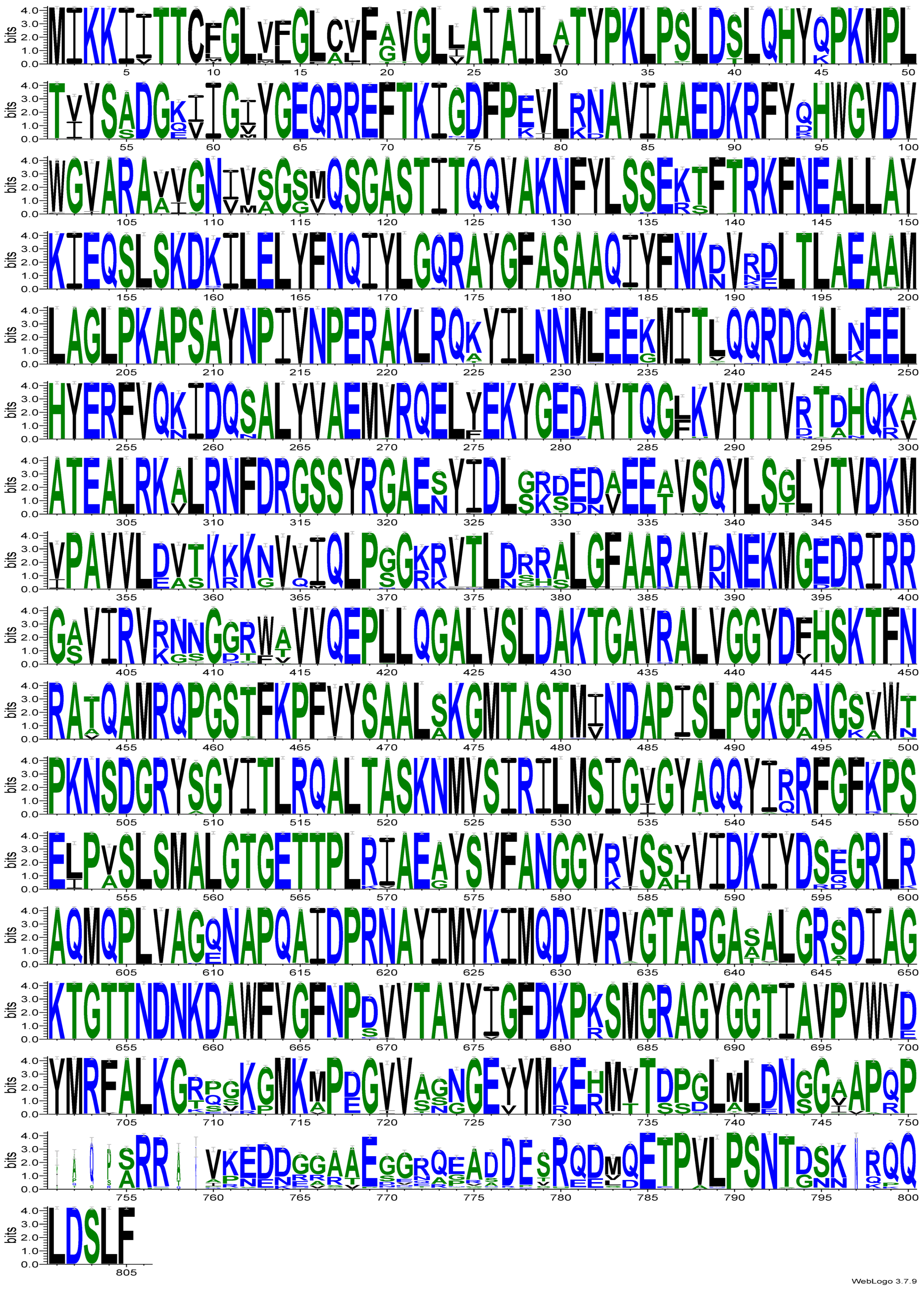

**FIG S2** Penicillin MICs and prevalence of *ponA*$^{L421P}$, *ponA*$^{A375T}$, and *ponA*$^{Y537C}$ amongst gonococcal isolates. Phylogeny of 6,082 representative gonococcal genomes after recombination detection is mapped in the center. The inner ring indicates the penicillin MIC, S is susceptible (MIC ≤0.06 µg/mL), I is intermediate (MIC 0.12–0.25 µg/mL), R is resistant (MIC 0.5-8 µg/mL), and HL-R is high level resistant (MIC >= 8). The second ring represents whether the *blaTEM* plasmid is present (purple), or absent (light blue) in the isolate. The *blaTEM* plasmid confers high-level penicillin resistance, and isolates that encode this plasmid usually have penicillin MICs above 8ug/mL. The third ring represents whether the strain possess a leucine residue 421 *ponA* (grey), or if residue 421 is mutated to proline, *ponA*$^{L421P}$ (black). The fourth ring represents whether the strain possess an alanine at *ponA* residue 375 (light pink), or if residue 375 is mutated to a threonine (magenta).The outer-most ring represents whether the strain possess a tyrosine at *ponA* residue 537 (light blue), or if residue 537 is mutated to a cystine (purple).

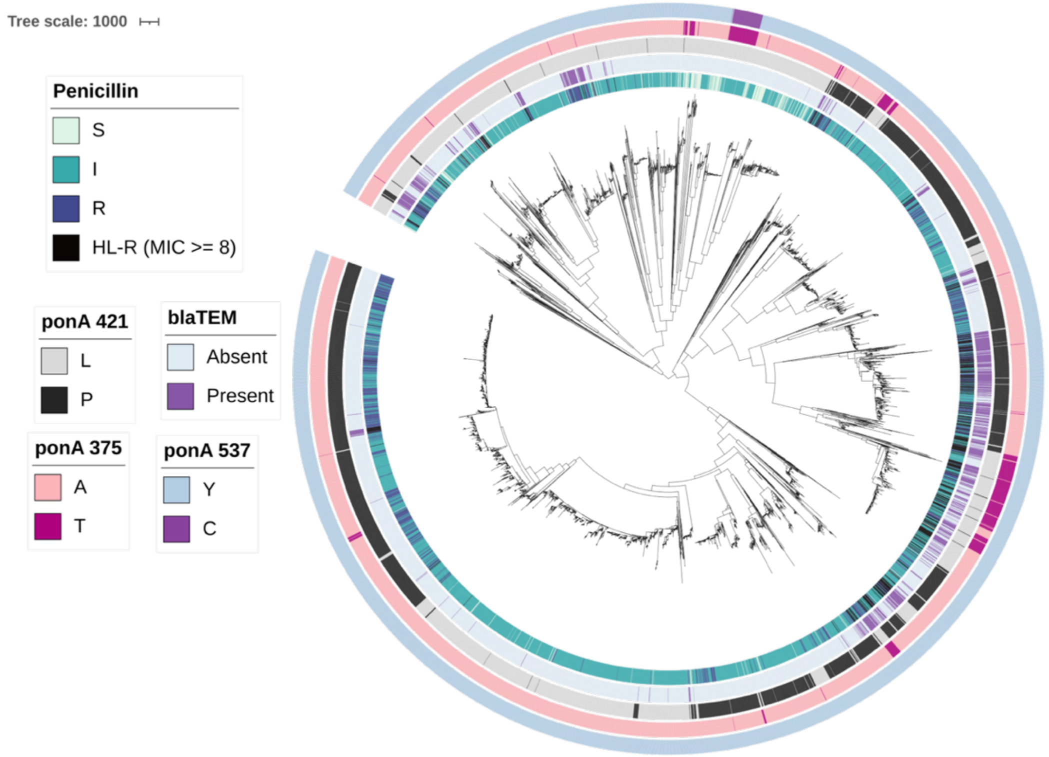

**FIG. S3** Relative read counts for each FA6140 *ponA*$^{L421X}$ *s*train in the inoculum cultures used in Trials 2 and 3**.** Percentage of reads, relative to the total number of reads, for each mutant in the starting inoculum from Trial 2 (A) and Trial 3 (B). Each trial represents a separate Illumina sequencing run, (see Fig. 6A for data from Trial 1).

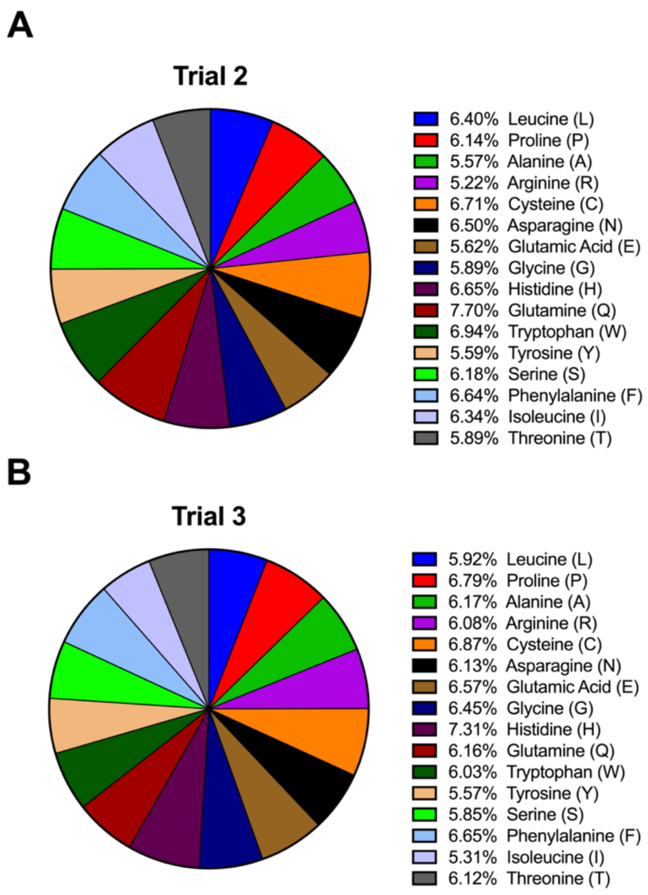

**FIG. S4** (A) Optical densities and (B) Illumina read counts for the cultures supplemented with 0.25 µg/mL penicillin. (B) Reads for each mutant in the inoculum were normalized to 1, as indicated by the dotted line at y=1. Bars represent read counts for each mutant at hour 36, relative to their inoculum value. Experiment was preformed 3 separate times (trials) and each trial represents a separate Illumina sequencing run; Trial 1 is shown in red, trial 2 in yellow, and trial 3 in blue.

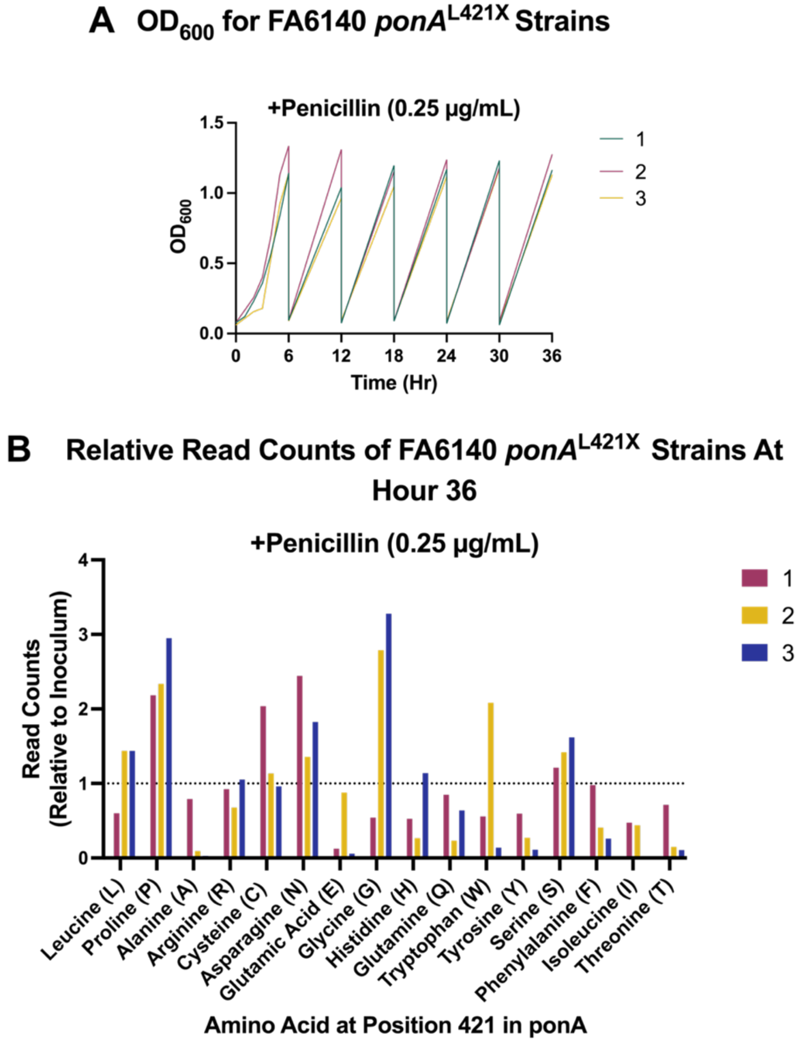

**FIG. S5** Illumina read counts for culture supplemented with 1.0 µg/mL penicillin in trial 2. Reads for each mutant in the inoculum were normalized to 1, as indicated by the dotted line at y=1. Bars represent read counts for each mutant at hour 36, relative to their inoculum value. Only hours 0-18 are shown, as cells did not survive passaging after hour 18. See Fig. 7 for data from trials 1 and 3.

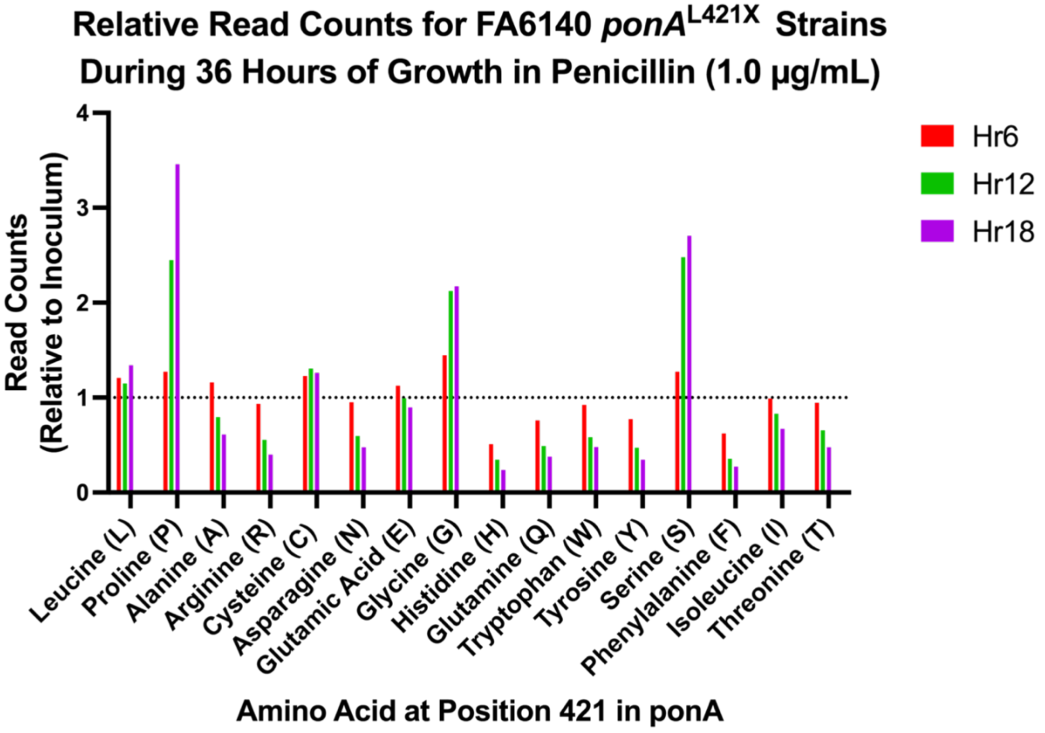
